# Proteomics Identifies Osteomodulin as a Promoter of Breast Cancer Bone Metastasis via CDK1 Activation

**DOI:** 10.1101/2023.11.03.565489

**Authors:** Sara Cabral, Joseph Parsons, Hannah Harrison, Thomas Kedward, Paul Fullwood, Katherine Spence, Diane V. Lefley, Danielle Barden, Jennifer Ferguson, Joanne Watson, Kalliopi Tsafou, Caron Behan, Mark J. Dunning, Nisha Ali, Balázs Győrffy, Janet Brown, Michael P. Smith, Penelope D. Ottewell, Ciara S. O’Brien, Chiara Francavilla, Robert B. Clarke

**Affiliations:** Division of Cancer Sciences, School of Medical Sciences, Faculty of Biology Medicine and Health, University of Manchester, Manchester UK; Division of Cellular and Molecular Function, School of Biological Sciences, Faculty of Biology Medicine and Health, University of Manchester, Manchester, UK; Division of Clinical Medicine, Medical School, University of Sheffield, Sheffield, UK; Stanley Center for Psychiatric Research, Broad Institute of MIT and Harvard, Cambridge, MA, USA; Histology Cancer Research UK Manchester Institute, Manchester, UK; Sheffield Bioinformatics Core, School of Medicine and Population Health, University of Sheffield, UK; Manchester University Foundation Trust, Wythenshawe Hospital, Manchester, UK; Manchester Breast Centre, University of Manchester, Manchester, UK; Department of Bioinformatics, Semmelweis University, H-1094, Budapest, Hungary; Department of Biophysics, Medical School, University of Pecs, H-7624, Pecs, Hungary; Institute of Molecular Life Sciences, HUN-REN Research Centre for Natural Sciences, H-1117, Budapest, Hungary; Department of Medical Oncology, Christie Hospital, Manchester, UK; DTU Bioengineering, Danish Technical University, Lyndby, Denmark

**Keywords:** breast cancer, stroma, bone metastasis, proteomics, phosphoproteomics, osteomodulin, CDK1, CDK1 inhibitor, cancer-associated fibroblasts

## Abstract

Metastasis to different organs remains the main cause of mortality in breast cancer. Molecular predictors of metastasis are limited as well as therapeutic options. Here, we conducted quantitative proteomics and phosphoproteomics analysis of patient-derived tumours, identifying osteomodulin (OMD) as a dysregulated protein and associated with bone metastases. Cancer-associated fibroblasts secrete OMD which increases breast cancer migration *in vitro* and promotes the formation of bone metastases *in vivo*. Downstream of OMD, phosphoproteomics identified the activation of cyclin-dependent kinase 1 (CDK1). The OMD-CDK1 signalling axis drives a pro-migratory and pro-survival phenotype *in vitro* and bone metastasis *in vivo*. Our findings highlight the importance of OMD and CDK1 in breast cancer bone metastasis and proposes an alternative therapeutic avenue for the treatment and the prevention of organ-specific metastases.

## Introduction

Metastases are responsible for 90% of breast cancer deaths ^1^ and are most commonly found in the bone, brain, liver and lungs of breast cancer patients ^2^. Bone is the most prevalent site of metastasis, representing around 70% of metastatic breast cancers cases^3^. Bone metastases are characterized by being mainly osteolytic, leading to debilitating symptoms such as pathological fractures, severe bone pain, and hypercalcemia^4^. Currently, there are no clinically available biomarkers nor targeted drug therapies against organ-specific metastases in breast despite the identification of different drivers of metastatic disease^3^. Radiologically detected, or symptomatic bone metastases are typically managed with bisphosphonates and receptor activator of NF-κB ligand (RANKL) inhibitors to alleviate symptoms and reduce bone osteolysis^4^. However, there is still the need to uncover more specific biomarkers and actionable combinations of therapeutic targets for patients at risk of developing metastases. Preventing bone metastasis or targeting bone organotropic drivers would greatly benefit patients given the capacity of the bone microenvironment to condition tumour cells to colonise other organs ^5–7^.

Omics technologies have the capacity to provide novel biomarkers and therapeutic targets from a variety of tissues, including patient-derived samples ^8^. For instance, genomic analysis of metastatic breast cancer patient samples or matched primary and metastatic breast cancer patient samples identified several potential therapeutic targets whose mutation was enriched in metastases ^9^. Transcriptomic analysis of breast cancer primary and metastatic samples also identified several transcripts dysregulated in breast cancer bone metastases compared to other metastatic sites ^10^. In contrast to genomics and transcriptomics, proteomics and phosphoproteomics have so far analysed primary breast cancer but not metastatic samples ^11^, for example identifying dysregulation in tumour cells metabolism ^12^, novel kinases in triple negative breast cancer patients ^13^, and signalling pathways associated with Epidermal Growth Factor Receptor 2 (HER2)-targeted therapy resistance ^14^.

Here, we set out to identify potential therapeutic targets and their biomarkers in breast cancer metastasis organotropism. Fresh frozen primary breast cancer resection samples that were clinically annotated as to the presence or absence of organ-specific metastases within 5 years of diagnosis (distant disease-free survival at 5 years) were analysed. We combined mass spectrometry-based proteomics and phosphoproteomics, bioinformatics, and functional assays *in vitro* and *in vivo*. This experimental pipeline identified Osteomodulin (OMD), a secreted member of the small leucine-rich proteoglycan family typically expressed by osteoblasts and associated with biomineralization processes^15,16^. We demonstrated that OMD is expressed and secreted by cancer-associated fibroblasts (CAFs) and stimulates the migration of estrogen receptor-positive (ER+) and ER-negative breast cancer cells. Using further phosphoproteomic analysis of OMD-stimulated breast cancer cells, we identified activation of Cyclin-dependent kinase (CDK1) to be necessary for BC migration, viability and bone metastasis. High OMD and CDK1 co-expression predicts metastatic outcome independently of other clinical and prognostic factors in a large cohort of breast cancer with over 15 years follow-up. Overall, this research establishes CAF-derived OMD and downstream CDK1 signalling as important regulator of breast cancer metastasis to the bone.

## Results

### Proteomic Analysis of Primary Breast Cancer Samples Identifies Osteomodulin (OMD) in Patients Developing Bone Metastases

To identify changes in proteins and phosphorylated proteins underlying breast cancer metastasis, we analysed a cohort of 39 primary breast cancer patient samples, of which 21 developed metastases within 5 years of diagnosis (here, referred to as metastatic) while 18 were distant disease free at 5 years (here, referred to as non-metastatic), by label-free mass spectrometry-based proteomics and phosphoproteomics (Figure 1A, Tables S1-3). Of the 39 patient samples, 21 were from patients who had hormone receptor positive (HR+) tumours, 9 who had human epidermal growth factor receptor 2 positive (HER2+) tumours and 9 who had triple negative (TN) tumours. Our cohort included patients who developed metastases at all four of the main metastatic sites in breast cancer at 5 years post-diagnosis (bone, brain, liver and lung) ^2^, thus providing a unique dataset (Figure 1A, Table S1). We identified and quantified a total of 6243 proteins (5106 in non-metastatic samples and 5741 in metastatic samples) and 14281 phosphorylated sites (11148 in non-metastatic samples and 13186 in metastatic samples) (Figure S1A-B, Tables S2-3). Pearson correlation for technical replicates in the proteomics data was above 0.95 on average and was above 0.6 on average in the phosphoproteomics data (Figure S1C-D), thus showing high correlation. Across phosphorylated sites, 62.8% were singly phosphorylated, 29.0% were doubly phosphorylated and 7.2% were more than doubly phosphorylated and the residue distribution across serine, threonine and tyrosine was 88.24%, 10.92%, 0.83%, respectively with a median score for phosphorylated peptides of 172 (Figure S1E-G). Therefore, the quality of our proteomics and phosphoproteomics analysis was consistent with previous publications ^11,17^.

**Figure 1.**
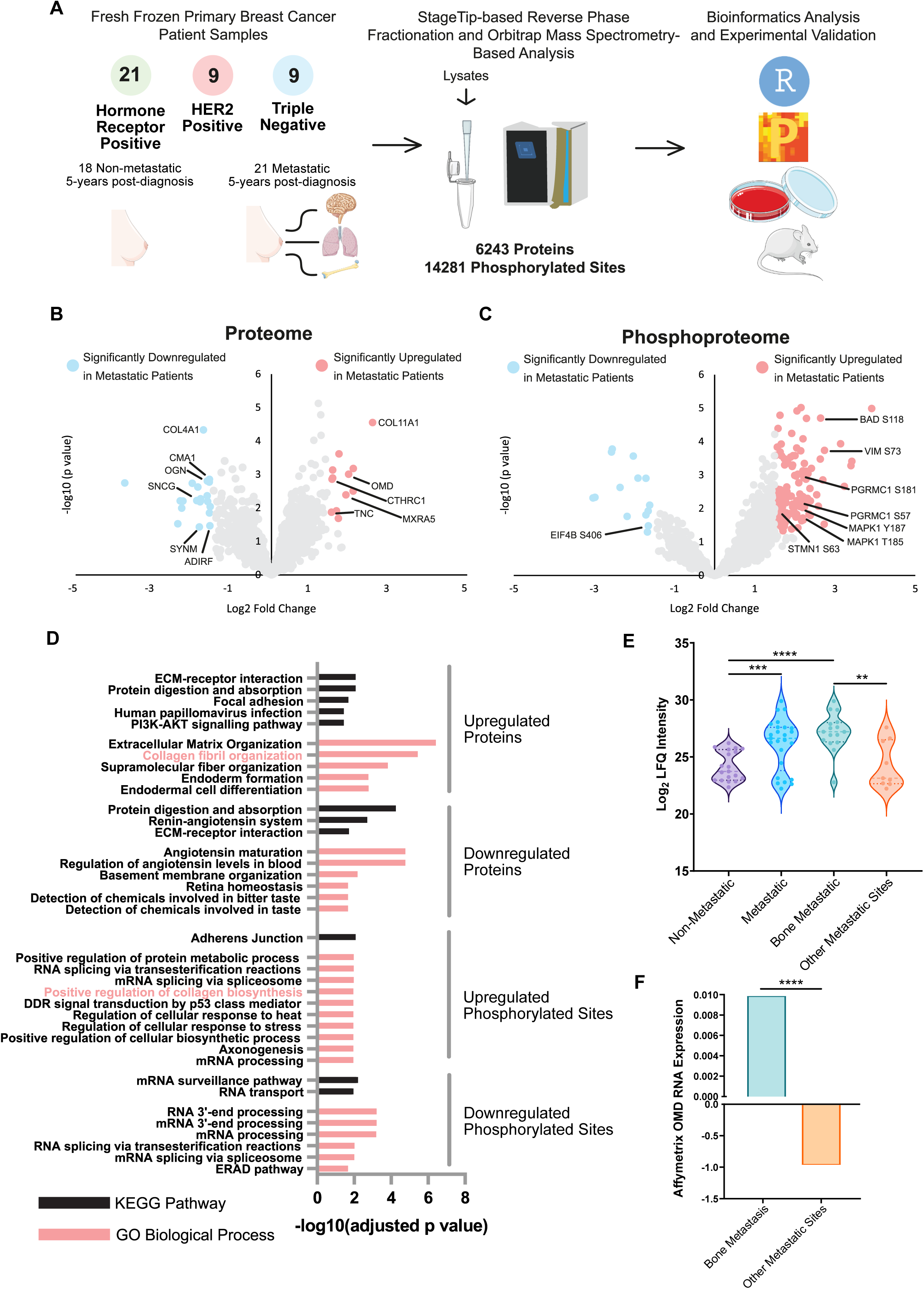
Proteomics analysis identifies enrichment of OMD in samples from patients who developed bone metastases **A,** Workflow of the proteomics and phosphoproteomics analysis of patient-derived samples. The characteristics of samples and the numbers of proteins and phosphorylated sites identified are indicated. Tissue and mouse images were downloaded from SmartServier Medical Art. **B, C,** Volcano plots showing differentially expressed proteins (B) or phosphorylated sites (C) based on Limma *p* value <0.05, Log2 Fold change >1.5 between metastatic and non-metastatic samples. Proteins or sites upregulated in patients who developed metastases are in orange while proteins and sites downregulated in patients who developed metastases are in blue. Proteins and phosphorylated sites previously associated with breast cancer metastasis regulation are labelled (see Table S7). **D,** KEGG and GO terms enrichment analysis of differentially regulated proteins and phosphorylated sites between metastatic and non-metastatic samples from B and C. Adjusted *p* values are generated using Fisher’s exact test with Benjamini-Hochberg correction. **E**, Osteomodulin (OMD) log2 normalised protein intensity in non-metastatic patients, metastatic patients, patients who developed bone metastases or patients who developed metastases at any other sites not including bone. Data represent mean +/− SEM of 15 non-metastatic samples compared to 20 metastatic samples, and 12 bone-metastatic samples compared to 8 non-bone metastatic samples (Limma *p* values ** <0.01, *** <0.001, **** <0.0001) and are visualised in the same graph. **F,** OMD mRNA levels in bone metastasis and in metastases to other organs from the GEO datasets (https://www.ncbi.nlm.nih.gov/pmc/articles/PMC6424882/). Data (37 bone metastatic patient samples, 148 other metastatic site patient samples) are represented as mean (Limma *p* values with Benjamini-Hochberg False Discovery Rate Correction < 0.0001****). See also Figures S1-2, Tables S1-7.

We focused on proteins and phosphorylated sites present in at least 80% of either non-metastatic or metastatic samples for further analysis (Tables S4-5). Differential expression analysis of patients who developed or did not develop metastases identified 33 differentially regulated proteins and 156 differentially regulated phosphorylated sites on 122 proteins (*p* value <0.05, Log2 Fold Change in intensity >1.5) (Figure 1B-C, Tables S4-5). To determine whether these differentially regulated proteins and phosphorylated sites had previously been associated with breast cancer metastasis, we compared our list of 33 dysregulated proteins and 156 dysregulated phosphorylated sites with an in-house curated database of metastasis-associated genes and proteins obtained by searching PubMed with the term “Breast Cancer Metastasis” for a total of 2472 entries (Table S6). We found that ADIRF, CMA1, COL11A1, COL4A1, CTHRC1, MXRA5, OGN, OMD, SNCG, SYNM, TNC proteins and BAD, MAPK1 (2 sites), PGRMC1 (2 sites), STMN1, EIF4B and VIM phosphorylated sites have been previously associated with breast cancer metastasis or metastasis-promoting phenotypes (Figure 1B-C, Tables S4-6). Thus, our proteomics and phosphoproteomics analysis identified both known and novel drivers of breast cancer metastasis.

Next, KEGG and GO enrichment analysis identified several biological processes among all the upregulated proteins and phosphoproteins, including collagen remodelling, previously associated with breast cancer ^18^ (Figure 1D, Table S7). Osteomodulin (OMD), a member of the small leucine-rich repeat protein family^15,16^, was among the upregulated proteins and phosphorylated sites associated with collagen synthesis and remodelling. Analysis of published datasets had previously found OMD gene expression to be correlated with breast cancer bone metastasis^10^. In our study, OMD protein was found to be significantly higher in patients who developed bone metastasis compared to patients who developed metastases in other sites (*p* value = 0.0012), or that did not develop metastases (*p* value = <0.0001) (Figure 1E, Tables S1, S4). Immunohistochemical analysis of the primary tumour from the same cohort of breast cancer samples detected OMD in 17 samples and showed heterogeneous staining in the tumour and stromal compartments (Figure S2). Consistent with OMD protein expression being associated with bone metastasis, we found that OMD mRNA was significantly higher in patients developing bone metastases compared to other metastatic sites based on three published breast cancer GEO datasets (Figure 1F) ^10,19^.

Overall, our proteomic and immunohistochemical analysis of primary breast cancer samples indicates that OMD expression is higher in breast cancer patients developing bone metastasis.

### OMD is Expressed by Cancer-Associated Fibroblasts and Regulates Cell Migration in HR+ and HR-Breast Cancer

As OMD was found in the tumour and in its microenvironment (Figure S2), we first tested whether OMD affected either cells in the tumour microenvironment or the epithelial tumour cells themselves. We used cell models of the primary and secondary (bone) tumour microenvironment including normal and cancer-associated fibroblasts (CAFs) (HMFU19 and 544R, respectively) ^20,21^, osteoblasts (HFOB 1.19) ^22^, and osteoclasts (RAW 264.7, post-differentiation with RANKL) ^23^. As a model of tumour epithelial cells, we used the epithelial breast cancer cell line MDA-MB-231 infected with lentivirus expressing either mApple as a control (231-mA) or an OMD-T2A linker-mApple construct (231-OMD) (Figure 2A-B). OMD expression was confirmed by immunoblotting in cell lysates from both starvation and complete media and in conditioned medium (CM) (Figure 2A-B), thus confirming that OMD is a secreted protein ^24^. We tested whether there were any effects on osteogenesis, a key process in the differentiation and turnover of bone tissues involved in sclerotic metastases ^25^, by assessing Alkaline Phosphatase Activity and calcium deposition via Alizarin Red staining. To mimic secreted OMD, we used recombinant OMD (rOMD) at increasing concentrations from 10ng/ml to 1µg/ml, which recapitulates the concentrations used by others^26^. Both osteogenesis and calcium deposition were significantly induced in the osteoblast cells upon differentiation (Figure S3A-C). However, compared to differentiated osteoblasts, alkaline phosphatase activity is unaffected by conditioned medium (CM) from 231-mA or 231-OMD or by increasing concentrations of rOMD, except for the highest concentration of rOMD which induces calcium deposition. Additionally, in the bone-resorbing osteoclasts, no significant induction of activity was seen upon treatment of osteoclasts with either CM from 231-mA and from 231-OMD cells or rOMD was observed, as the number and size of pores was consistent across all conditions (Figure 2A-B, Figure S3D-G). Collagen contractility was also assessed to understand whether OMD affects matrix remodelling in normal fibroblasts, CAFs and differentiated osteoblasts. rOMD and CM derived from 231-OMD and 231-mA cells had no effect on the collagen remodelling capabilities of each of the tested cells (Figure S3H-J).

**Figure 2.**
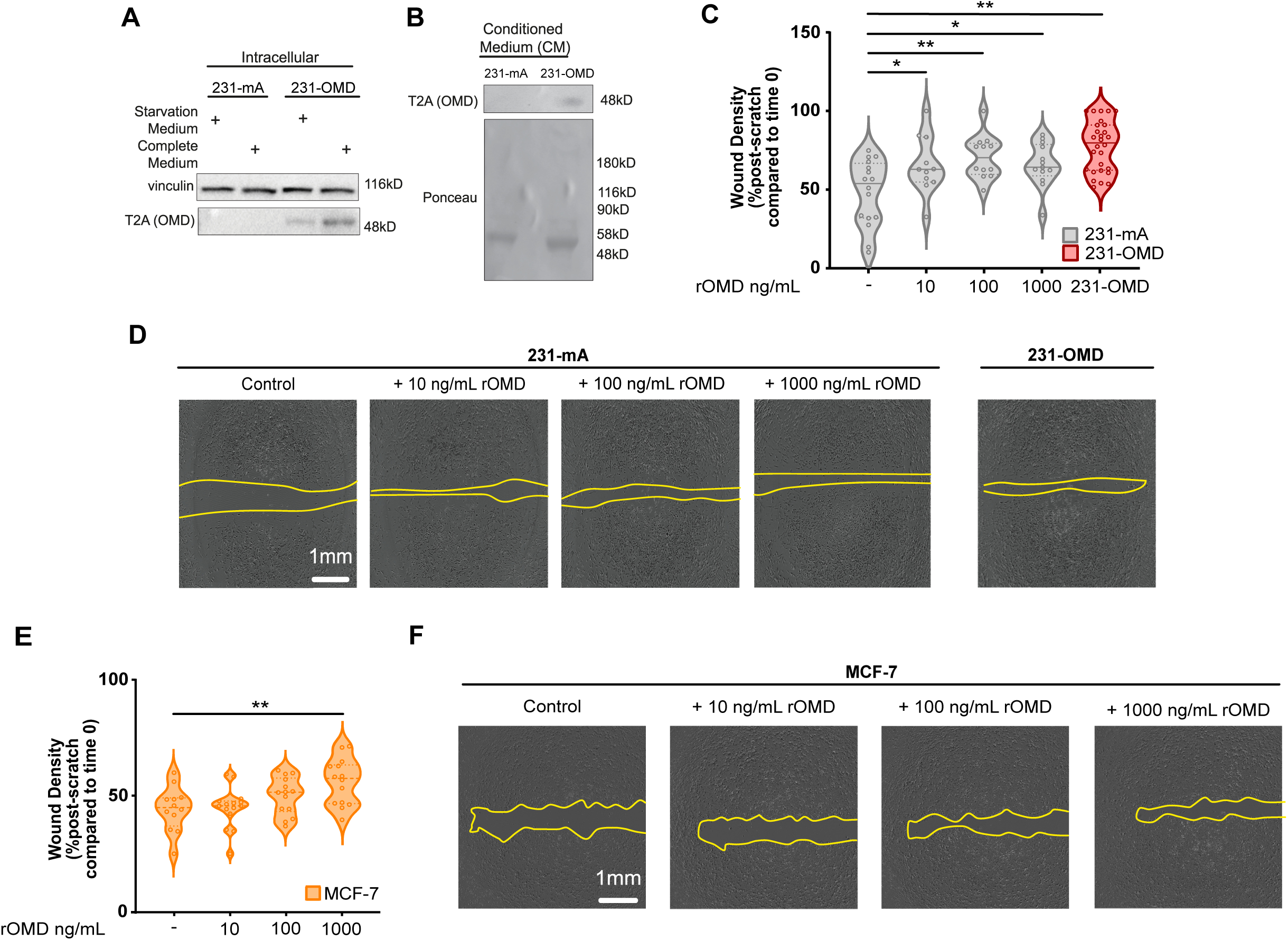
OMD regulates cell migration *in vitro* in both ER+ and ER-cell line models of breast cancer. **A, B,** Immunoblotting analysis with the indicated antibodies of total lysates from MDA-MB-231 cells stably transfected with mApple (231-mA) or an OMD-T2A-mApple construct (231-OMD) and grown in complete or starved medium (A) and of conditioned medium from the same cells (B) N=3 independent biological replicates. **C,** Cell migration of 231-mA (grey), 231-OMD (red), 231-mA treated with 10, 100 or 1000 ng/ml of rOMD assessed by wound density (percentage of the wound filled with cells 48hrs post-scratch compared to time zero). **D,** Representative images from C. Scale bar = 1mm. **E,** Cell migration of MCF-7 (orange) treated with 10, 100 or 1000 ng/ml of rOMD assessed by wound density (percentage of the wound filled with cells 48hrs post-scratch compared to time zero). **F,** Representative images from E. Scale bar = 1mm. **C, E,** Data represents the mean +/− SEM of N = 3 independent biological replicates, each including at least 3 technical triplicates. *p* =< 0.05*, < 0.01**, < 0.001***, < 0.0001**** (One Way ANOVA with Dunnett’s Multiple Comparison Test, compared to untreated control). **D, F,** The yellow lines indicate wound gap at 48h. See also Figures S3-4.

Using 231-OMD cells, we next tested whether OMD expression affected the behaviour of breast cancer cells. We assessed behaviour by looking at effects on extracellular matrix remodelling, proliferation and stemness of breast cancer cells by conducting collagen contractility assays, measuring EdU incorporation, and mammosphere formation and self-renewal assays, respectively. We found that OMD overexpression had no effects on any of these behaviours (Figure S4A-D). We next assessed the effects of OMD on cell migration using a scratch wound assay and cell invasion using a Boyden chamber assay. Although invasion was not significantly altered between 231-mA and 231-OMD cells in our experimental conditions, OMD expression significantly increased cell migration (Figure 3C-D, Figure S4E). Additionally, treatment with increasing concentrations of rOMD significantly induced 231-mA cell migration in a dose-dependent manner (Figure 3C-D). We also tested the effect of rOMD on the migration of ER+ MCF-7 cell line (Fig 3F-G), and confirmed that the highest concentration of rOMD increased MCF-7 migration. However, OMD expression did not affect cell motility of the bone-tropic MDA-MB-231 (231-Bo) cell line ^27^. Indeed, cell migration of the bone-tropic MDA-MB-231 cell line expressing OMD (231-Bo OMD) was similar to control cells (231-Bo) (Figure S4F-I).

**Figure 3.**
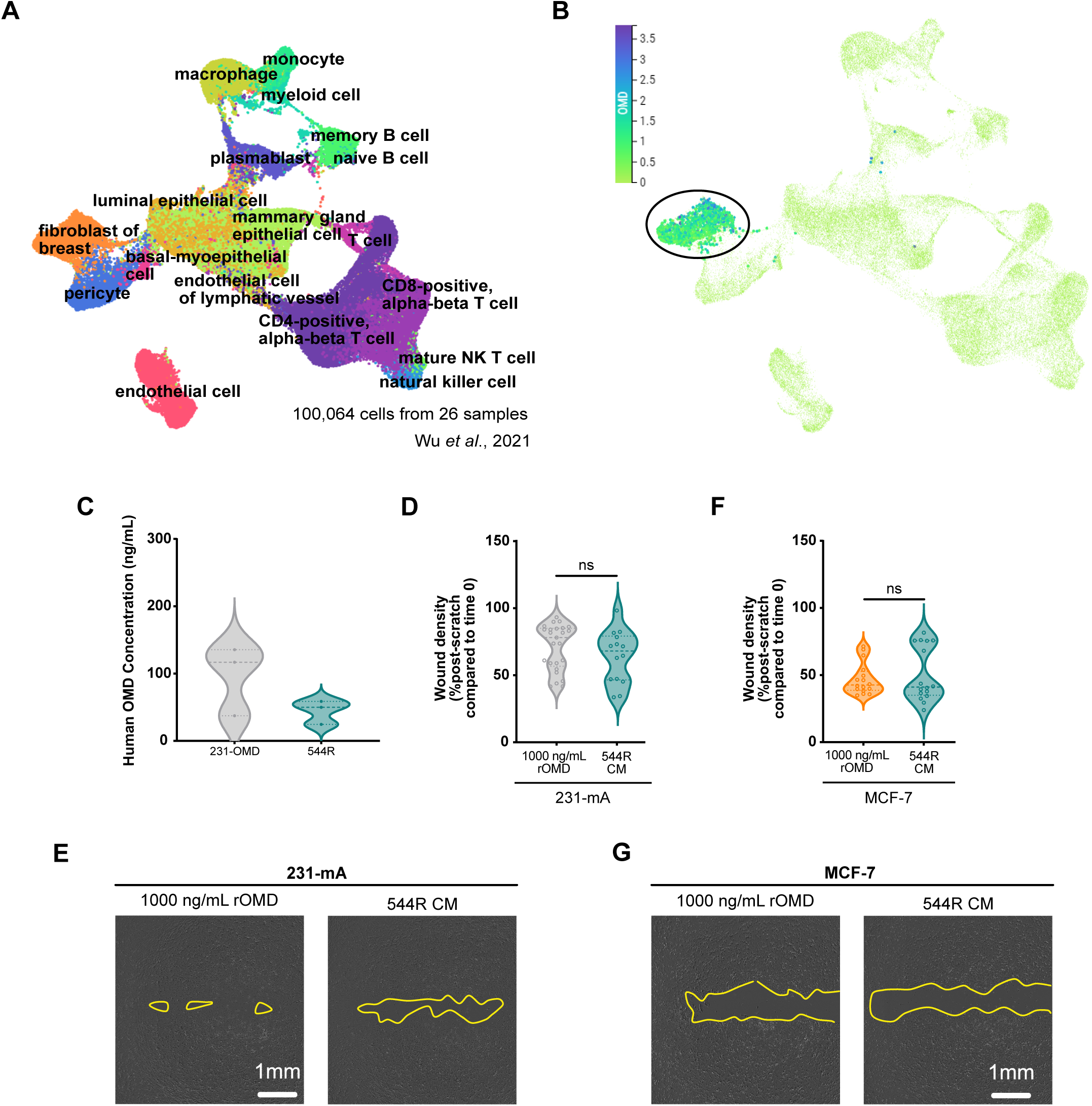
OMD is expressed by cancer-associated fibroblasts in patient samples and fibroblast-derived conditioned media drives cell migration. **A,** Single cell RNAseq UMAP analysis of 26 primary breast cancer samples showcasing all the cell types^78^. **B,** UMAP analysis of OMD RNA levels from the dataset in A. UMAP plot of OMD expression across all cell types analysed using CELLxGENE^28^. Black circle highlights the fibroblast population as containing cells expressing OMD. **C,** ELISA-based quantification of OMD secretion from conditioned media (CM) collected from 231-OMD and 544R cells. Data represents the mean +/− SEM of N = 3 independent biological replicates, each including 2 technical replicates. **D,** Cell migration of 231-mA cells treated with 1000 ng/mL rOMD or with CM derived from 544R (blue) assessed by wound density (percentage of the wound filled with cells 48hrs post-scratch compared to time zero). **E,** Representative images from D. Scale bar = 1mm. **F,** Cell migration of MCF-7 treated with 1000 ng/mL rOMD or with CM derived from 544R (blue) assessed by wound density (percentage of the wound filled with cells 48hrs post-scratch compared to time zero). **G,** Representative images from F. Scale bar = 1mm. **D, F,** Data represents the mean +/− SEM of N = 3 independent biological replicates, each including at least 3 technical triplicates (Mann-Whitney Test). **E, G,** The yellow lines indicate wound gap at 48hrs. See also Figure S5.

Based on the pattern of OMD staining in our cohort (Figure S2) and to determine which cells in the tumour microenvironment were expressing OMD, we analysed available breast cancer single cell RNAseq expression data including both stromal and epithelial cells^28^. We found that CAFs but not other stromal cells consistently expressed OMD, and not epithelial cells as previously hypothesized (Figure 3A-B). To further confirm OMD expression in the stroma, we stained a tissue microarray of 482 breast cancer patient samples^29^. OMD staining was detected only in stromal areas and not in tumour epithelium (Figure S5). The single cell RNAseq and immunostaining analyses establish that OMD gene and protein are expressed in CAFs. To validate this, we proceeded to collect CAF CM from our selected CAF cell line 544R. Using ELISA, we showed that CAF CM contained 45 ng/mL OMD while the 231-OMD CM contained 96 ng/mL (Figure 3C). The CAF CM stimulated both 231-mA and MCF-7 cell migration similarly to what we observed by adding rOMD (Figure 3D-G) and in line with previous results (Figure 2). Overall, we established that OMD increases cell migration in ER+ and ER-breast cancer and is secreted in the tumour microenvironment by CAFs, suggesting that fibroblasts play a role in inducing breast cancer metastasis to the bone.

### Phosphoproteomics Identifies an OMD-CDK1 Axis Driving Cell Migration and Viability

To ascertain the mechanism underlying the role of OMD in breast cancer cell migration, we analysed changes in the proteome and phosphoproteome of 231-OMD cells compared to 231-mA (Figure 4, Figure S6). 231-Bo and 231-Bo OMD cells were used as further controls. We identified and quantified 3971 proteins and 9423 phosphorylated sites (Tables S8-9). Pearson correlation between biological replicates of the proteomic and phosphoproteomic data were consistently over 0.97 and 0.6 respectively (Figure S6B-C). Across phosphorylated peptides, 41.9% were singly phosphorylated, 47.6% were doubly phosphorylated, and 10.5% were more than doubly phosphorylated and the distribution of phosphorylated site residue across serine, threonine and tyrosine was 85%, 11.3% and 3.7%, respectively, indicating enrichment on tyrosine residues, with a median phosphorylated peptide score of 182 (Figure S6D-F). All the quality controls were in line with previous publications ^11,17^. Differential expression analysis showed no differences between proteins expressed by 231-OMD and 231-mA (P < 0.05, Log2 Fold change > 1, Table S10) but identified 53 distinct phosphorylation sites, of which 18 were upregulated and 35 were downregulated (Fig. 4A, Table S11). Of the 18 sites specifically phosphorylated at higher levels in 231-OMD compared to 231-mA cells, only FOS like 1 (FOSL1) phosphorylation on serine 26 has a known cancer-associated function according to Phosphosite plus ^30^ and has been shown to promote migration in MCF-7 breast cancer cells^31^. Of the 35 downregulated phosphorylated sites in 231-OMD compared to 231-mA cells, 5 were also downregulated in 231-Bo OMD compared to 231-Bo cells, suggesting that these sites are unlikely associated with the 231-OMD pro-migratory signalling mechanism and thus they were excluded from further analysis (Table S11). Of the remaining 30 phosphorylated sites specifically downregulated in 231-OMD compared to 231-mA cells, 6 had metastasis-associated functions, such as promoting cell migration, invasion, or drug resistance according to Phosphosite plus ^30^ (Figure 4A). Only one of these 30 downregulated sites was associated with inactivation of protein activity which was Tyrosine 15 (Y15) on the cell cycle regulator, Cyclin-dependent kinase 1 (CDK1, CDC2) ^32^ (Figure 4A, Table S11). We confirmed the downregulation of Y15 on CDK1 in 231-OMD compared to 231-mA, 231-Bo OMD, and 231-Bo cells by quantified immunoblotting (Figure 4B-C). Next, we searched for evidence of the OMD-CDK1 crosstalk in available phosphoproteomics datasets analysing breast cancer patient samples. We found an inverse relationship between OMD expression and reduction of CDK1 inhibitory phosphorylation in two independent phosphoproteomics datasets analysing 105 and 122 breast cancer patients, respectively ^11,33^. In these phosphoproteomics analyses, CDK1 inhibitory phosphorylation at Y15 ^32^ was significantly reduced in patients where OMD expression was above the mean compared to patients where OMD expression was below the mean (Figure 4D-E). Therefore, the OMD-CDK1 axis may regulate the function of OMD.

**Figure 4.**
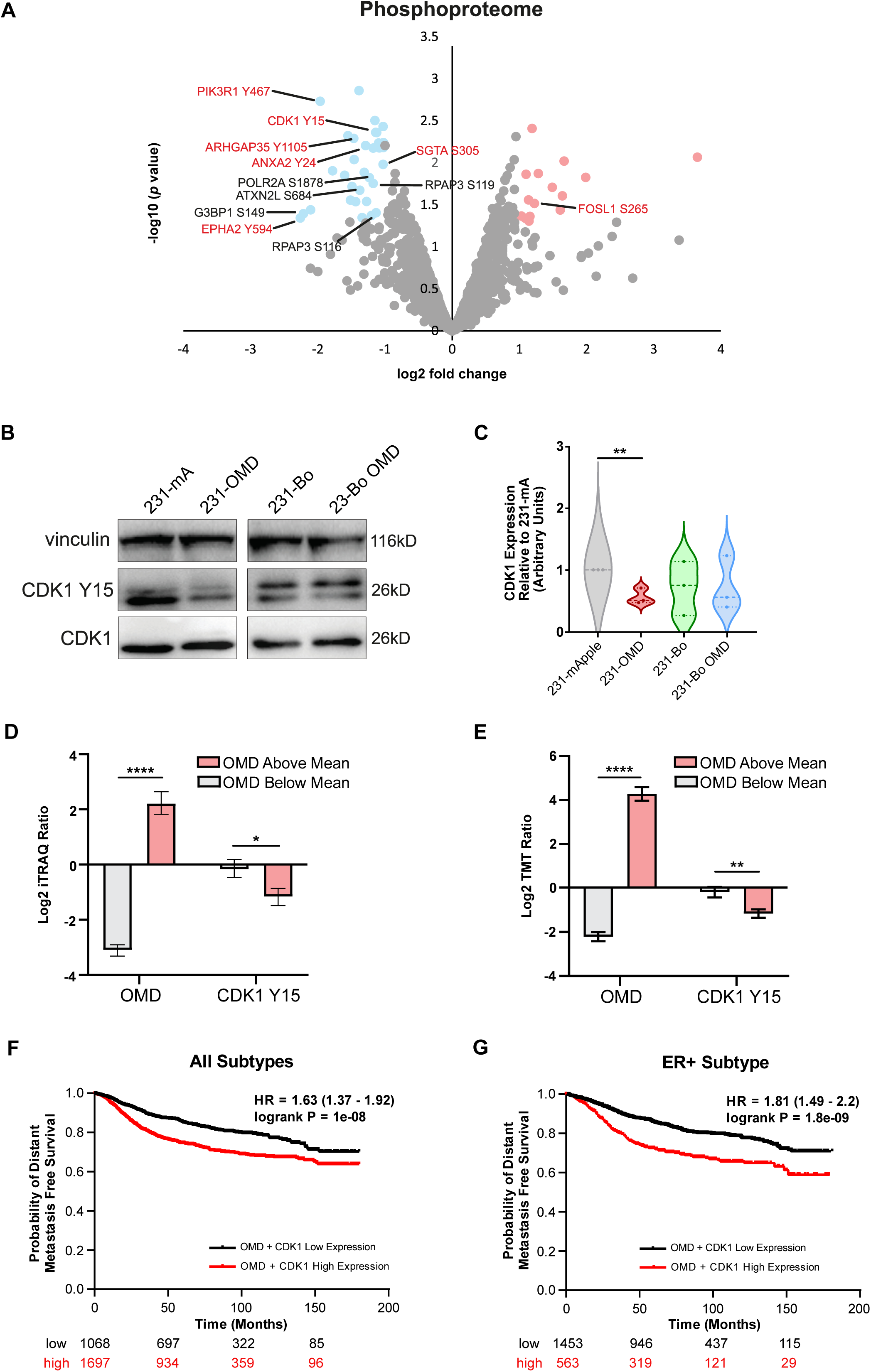
Phosphoproteomics identifies the OMD-CDK1 axis in breast cancer and patient-derived samples. **A,** Volcano plot showing differentially regulated phosphorylated sites in 231-OMD cells compared to 231-mApple cells based on Limma *p* value <0.05, Log2 Fold Change >1. Orange, upregulated in 231-OMD; blue, downregulated in 231-OMD; black text, downregulated sites in 231-OMD compared to 231-mApple and in 231-Bo OMD compared to 231-Bo; red text, sites with metastasis-associated functions according to Phosphosite plus. **B, C,** Immunoblotting analysis with indicated antibodies of 231-mApple, 231-OMD, 231-Bo and 231-Bo OMD cell lysates (B) followed by quantification via densitometry and normalisation to vinculin (C). Data is presented as the ratio between the intensity of CDK1 phosphorylated on Y15 and the intensity of the loading control. Values were calculated using ImageJ and are presented relative to 231-mA band intensity. N = 3 independent biological replicates p =< 0.05*, < 0.01**, < 0.001***, < 0.0001**** (Student’s t-test). **D, E,** Expression of OMD and of phosphorylated CDK1 on Y15 from two independent proteomics and phosphoproteomics datasets ^11,33^. Data is presented as mean +/− SEM of Log2 iTRAQ ratios (N = 83) and Log TMT ratios (N = 122), respectively, and compared to an internal standard. *p* =< 0.05*, < 0.01**, < 0.001***, < 0.0001**** (Student’s *t*-test). **F, G,** Kaplan-Meyer analysis of distant metastasis free survival with high and low expression of OMD and CDK1 in all breast cancer patients (F) and those with the estrogen receptor positive (ER+) subtype (G) for 180 months. See also Figures S6, Tables S6-12.

To investigate the contribution of OMD and CDK1 to breast cancer metastasis, we analysed 2,765 breast cancer cases from KM Plotter^34^. We investigated high expression (n = 1697) versus low expression (n = 1068) of both OMD and CDK1 using optimal cutoff over a follow-up period of 180 months. The hazard ratio (HR) of high versus low expression was 1.63, indicating a higher risk of recurrence in patients with high expression of OMD and CDK1 (Figure 4F). For ER+ patients, we investigated distant metastasis-free survival (DMFS) for high versus low expression and revealed a hazard ratio (HR) of 1.81, demonstrating a higher risk of relapse for patients with this subtype (Figure 4G). We further analysed the data to ensure that OMD and CDK1 were independent prognostic factors (Table S12). In the analysis of all tumors, the signature demonstrated independent prognostic value for DMFS, with a *p* value of 0.045. In ER+ tumors, the signature showed strong prognostic performance, with a *p* value of 1.1 × 10⁻⁴. It surpassed all clinical parameters, including HER2 expression (*p* = 0.055), lymph node status (*p* = 0.104), tumor grade (*p* = 0.025), tumor size (*p* = 0.0002), and patient age (*p* = 0.552). These findings underscore the independent prognostic impact of the signature, with its strongest predictive value observed in ER+ breast cancer cases (Figure 4G). Overall, the analysis found higher OMD/CDK1 to be a strong indicator of recurrence independent of other prognostic variables, especially in ER+ patients, 20-40% of whom experience metastasis and 70% of these correspond to bone.

### CDK1 Regulates Cell Migration and Viability *in vitro*

To verify the role of CDK1 downstream of OMD, we inhibited CDK1 using 10µM of the CDK1 inhibitor (CDK1i) RO-3306 ^35^ and assessed cell migration in the 231-mA and 231-OMD cells. As expected, 231-OMD cells migrate more than 231-mA cells. However, only in 231-OMD was cell migration significantly reduced upon treatment with 10µM RO-3306 (Figure 5A). To test whether this reduction in migration was specifically downstream of CDK1, we inhibited CDK4/6 with 10µM of the clinically relevant inhibitor Palbociclib (CDK4/6i) and showed that its inhibition did not significantly reduce 231-OMD cell migration (Figure 5A). Next, we investigated the ability of CDK1 inhibition to block rOMD or CAF-derived CM in both ER+ and ER-cells. We confirmed increased migration in response to rOMD which was effectively abrogated by RO-3306 in both 231-mA and MCF-7 cells (Figs. 5B-C). The increased migration observed after incubation with CAF-derived CM was also blocked by the CDK1i in both ER+ and ER-cell lines (Figure 5D). Therefore, CDK1 is specifically activated in breast epithelial cells to regulate cell migration *in vitro* and its inhibition decreases migration in response to OMD.

**Figure 5.**
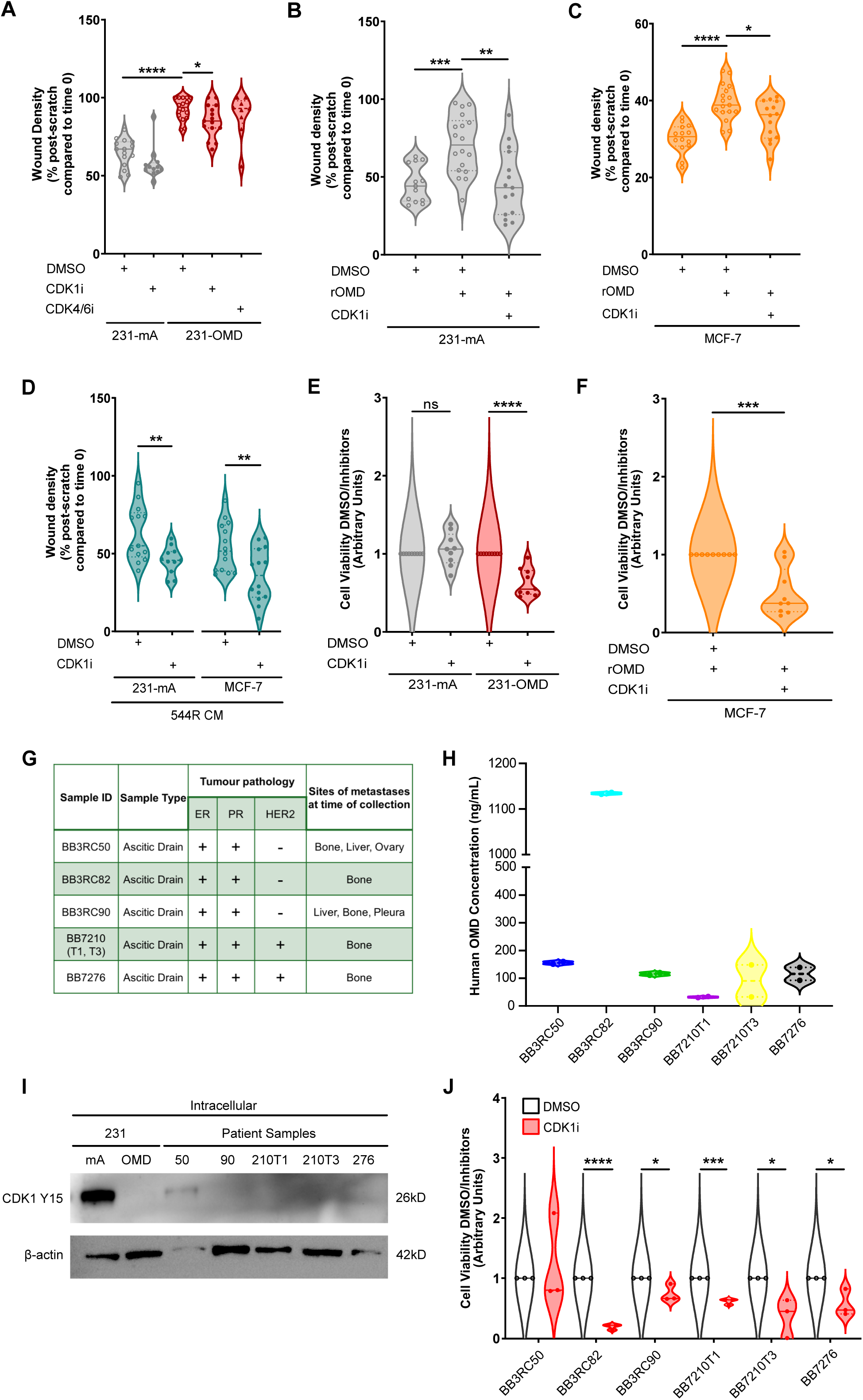
CDK1 regulates cell migration and viability downstream of OMD *in vitro*. **A,** Cell migration of 231-mA and 231-OMD treated with either DMSO, 10 µM RO-3306 (CDK1i) or 10 µM Palbociclib (CDK4/6i) assessed by wound closure and measured by wound density. **B, C,** Cell migration of 231-mA (B) and MCF-7 (C) cells treated with either DMSO or 1000 ng/mL rOMD in the presence or absence of 10 µM RO-3306 assessed by wound closure over 48hrs and measured by wound density. **D,** Cell migration of 231-mA and MCF-7 cells treated with the 544R conditioned medium (CM) in the presence or absence of 10 µM RO-3306. **E, F,** Relative viability of 231-mA (E) and MCF-7 (F) following treatment with 10 µM CDK1i. Data is represented as mean +/− SEM of N >= 3 independent biological replicates. *p* =< 0.05*, < 0.01**, < 0.001***, < 0.0001**** (Student’s *t*-test). **G,** Table showing the 5 patients from which metastatic effusions were collected along with their clinical characteristics and metastatic sites at time of collection. BB7210 had two drains collected at different timepoints (T1, T3). ER – Estrogen receptor, PR – Progesterone Receptor, HER2 – Human Epidermal Receptor 2. **H,** ELISA-based quantification of OMD in the metastatic fluid of the patients described in G. Data represents the mean +/− SEM of N = 1, each including two technical replicates. **I,** Immunoblotting analysis with the indicated antibodies of total lysates from fresh frozen samples derived from the patients illustrated in G, with the exception of BB3RC82. **J,** Relative viability of 6 metastatic fluid samples following treatment with 10 µM CDK1i. Data is represented as mean +/− SEM of N>= 1 independent biological replicates. *p* =< 0.05*, < 0.01**, < 0.001***, < 0.0001**** (Student’s *t*-test). **A, B, C,** Data represent the percentage of the wound filled with cells 48 hours post-scratch. N = 3 biologically independent replicates. p =< 0.05*, < 0.01**, < 0.001***, < 0.0001**** (One Way ANOVA with Dunnett’s Multiple Comparison Test, compared to untreated control). See also Figures S7, Tables S13-14.

Given the known role of CDK1 in regulating cell cycle and cell viability ^32^, we studied the effect of CDK1 inhibition on cell viability in both ER+ and ER-cells. We performed an Alamar blue assay ^36^ on 231-mA, 231-OMD, and MCF-7 cell models and found that CDK1 inhibition only reduced the viability of 231-OMD and MCF-7 cells treated with rOMD, but not 231-mA (Figs. 5E-F).

Finally, to test the clinical relevance of the OMD-CDK1 signalling axis, we used patient breast cancer cells derived from metastatic pleural effusions and ascitic fluids (Figure 5G, Figure S7A). Firstly, we assessed the effect of CDK1 activity on the viability of 5 metastatic patient-derived breast cancers *in vitro*, of which 4 showed decreased cell viability in response to CDK1 inhibition (Figure S7B). Phosphoproteomics and proteomics analysis of these samples produced a high-quality dataset (Figure S7C-G, Tables S13-14) and identified the downregulation of Y15 on CDK1 in the 4 samples that responded to CDK1 inhibition and reduced CDK1 Y15 phosphorylation significantly correlated with reduced viability in response to 10µM RO-3306 treatment (Figure S7B, S7H, Table S14). Next, we studied 6 ER/PR positive samples from ascites that had bone metastasis at the time of collection, 3 of which had amplification of HER2 (Figure 5G). The ascitic fluid containing the breast cancer cells expressed OMD at concentrations ranging from 30 to 1100 ng/mL (Figure 5H). We tested CDK1 Y15 phosphorylation in the breast cancer cells derived from the ascitic fluid and demonstrated low or no detectable phosphorylation, suggesting high activity of CDK1^37^ (Figure 5I). We also assessed cell viability of these samples upon treatment with the CDK1 inhibitor RO-3306 which demonstrated that 5 out of 6 samples responded (Figure 5J). The sample that did not respond (BB3RC50) was the only sample where CDK1 Y15 phosphorylation was evident by immunoblotting (Figure 5I). These data highlight the crucial role of active CDK1 downstream of OMD in regulating the viability of patient-derived breast cancer samples, including those with bone metastases at the time of collection (Figure 5G).

Overall, these results indicate a mechanistic role for CDK1 downstream of OMD secreted from fibroblasts in cellular processes relevant to metastasis such as cell migration and cell survival, including in breast cancer cells derived from patients with bone metastases.

### OMD-CDK1 Axis Drives Bone Metastasis and Bone Colonization *in vivo*

Next, we assessed the effect of OMD on metastatic potential *in vivo*. To do this, we injected 231-mA, 231-OMD, and 231-BO (positive control for bone metastases) cells by intracardiac injection into four athymic nude mice (Figure 6Ai). We found no significant differences in the lung metastatic potential of 231-mA, 231-OMD, and 231-BO cells (Figure 6B). However, in femoral bone flushes, the number of cytokeratin 19 (CK19)-positive cells, which is a marker for human epithelial cells ^27^, was significantly higher in 231-OMD compared to 231-mA (Figure 6C-D). As expected, the bone-tropic 231-Bo also colonized the bone. This finding confirms a role for OMD in the colonization of bone by breast cancer cells *in vivo*.

**Figure 6.**
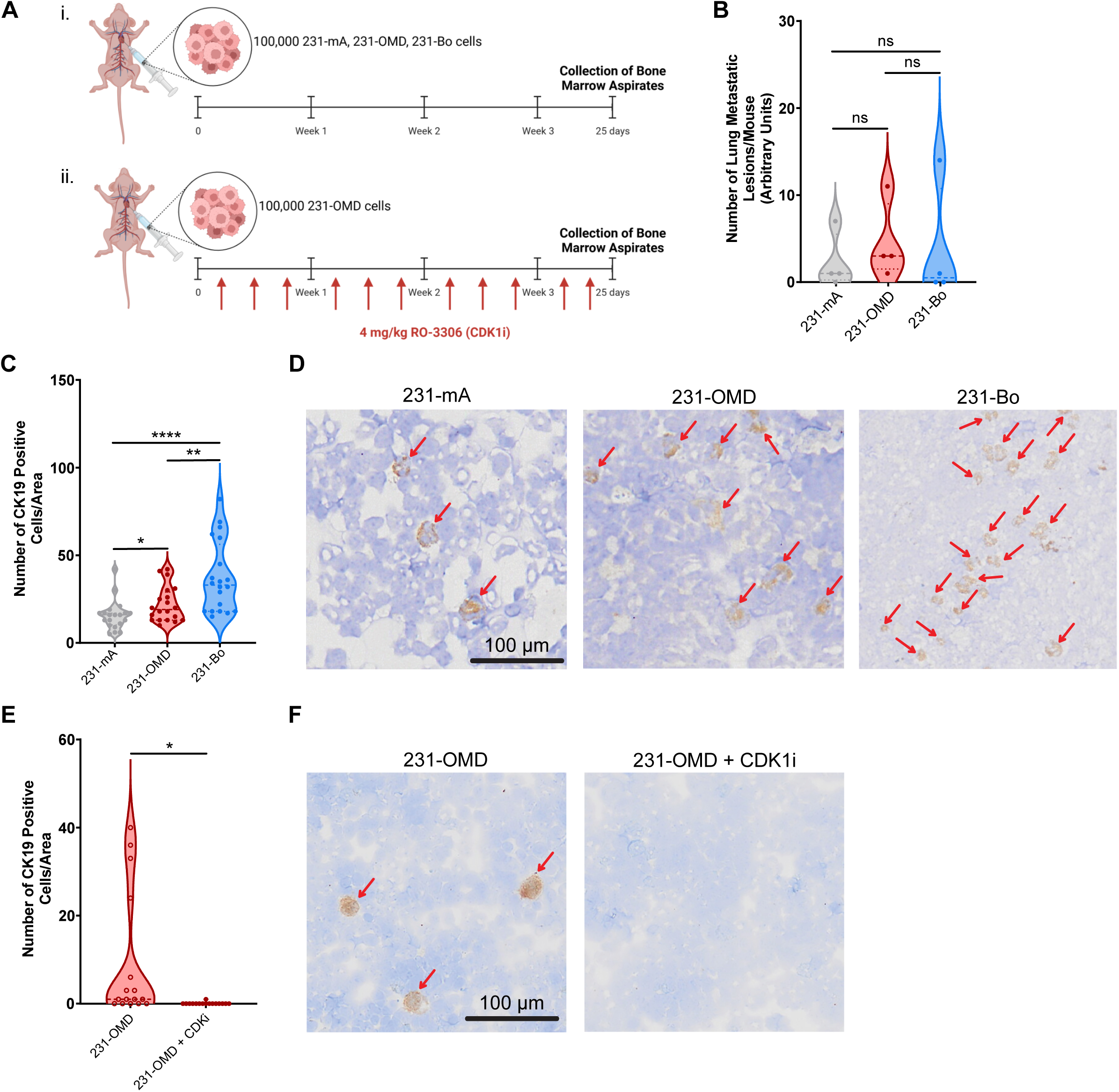
The OMD-CDK1 axis drives bone metastasis and bone colonization *in vivo*. **A,** Diagram illustrating the set-up of the *in vivo* experiments in athymic nude mice following the intracardiac injection of 100,000 231-mA, 231-OMD, or 231-Bo cells (i) and of 100,000 231-OMD cells followed by treatment with 4 mg/mL CDK1i (ii). Figure designed using Biorender.com. **B,** Number of metastatic lung lesions of over 100 µm in diameter (in at least 1 dimension) following intracardiac injection in N = 4 mice per cell line. **C,** Number of cytokeratin 19 (CK19)-positive human cells identified in cytospun bone flushes from the same mice. *p* =< 0.05*, < 0.01**, < 0.001***, < 0.0001**** (Mean +/− SEM, Student’s *t*-test, each compared to 231-mA). **D,** Representative images from C where red arrows indicated CK19-positive cells. **E,** Number of CK19-positive human cells identified in cytospun bone flushes following intracardiac injection and treatment with CDK1 inhibitor 4 mg/kg RO-3306 or vehicle control 3 times per week for 25 days in N = 4 mice per condition. *p* =< 0.05*, < 0.01**, < 0.001***, < 0.0001**** (Mean +/− SEM, Student’s *t*-test, each compared to 231-mA). **F,** Representative images from E where red arrows indicate CK19-positive cells.

Since CDK1 regulates cell migration and viability in 231-OMD cells, we next tested whether the OMD-CDK1 axis regulated bone colonization *in vivo.* To do this, we performed intracardiac injection of 231-OMD cells into four athymic nude mice and tested whether inhibition of CDK1 affected the formation of bone metastasis (Figure 6Aii). We detected numerous CK19-positive cells in bone flushes of mice treated with the vehicle control but almost no CK19-positive cells in those treated with the CDK1 inhibitor RO-3306 (Figure 6E-F). Altogether, these *in vivo* data indicate a role for the OMD-CDK1 axis in bone colonization by breast cancer cells.

In conclusion, we combined quantitative proteomics and phosphoproteomics from cancer patients with bioinformatics analysis and functional assays *in vitro* and *in vivo* and showed that OMD derived from CAFs regulates cell migration, cell viability and bone metastatic capabilities through a CDK1-dependent mechanism.

## Discussion

The yet unmet clinical need for novel biomarkers and targeted therapies to prevent and treat breast cancer metastasis is demonstrated by the still very high number of deaths from breast cancer each year of which 90% is caused by metastatic disease ^2^. Here, we combined quantitative proteomics and phosphoproteomics from breast cancer patient-derived samples and cell models with bioinformatics analysis and functional assays *in vitro* and *in vivo*. We demonstrated that OMD is expressed and secreted by cancer-associated fibroblasts (CAFs) and stimulates the migration of ER-positive and ER-negative breast cancer cells. We found that OMD requires the activation of CDK1 to regulate BC cell migration and cell viability and to promote bone metastasis *in vivo*.

Our study establishes that proteomics and phosphoproteomics of patient samples with known outcome at 5 years will identify novel drivers of metastasis. This time period was selected based on the available clinical outcome data. The aim was to derive proteins that could be used as biomarkers or drug targets in the clinic, as dysregulated genes or RNA transcripts identified in patient samples poorly correlate with the proteomics changes seen in metastases ^8,33^. Among the limitations of proteomics analysis of large cohort of patient samples are the availability of fresh frozen rather than paraffin-embedded (not yet suitable for phosphoproteomics) matched primary and metastatic samples and of the clinical information associated with them. Our proteomics and phosphoproteomics analysis of fresh frozen primary breast cancer resection samples provides therefore a unique resource as samples were manually annotated to determine the presence or absence of organ-specific metastases within five years of diagnosis ^38,39^. More recent approaches, including spatial proteomics^40^ and single-cell proteomics^41^ analysis of large cohort of clinical samples, will enable further discoveries in the near future.

Our proteomics analysis identified OMD, a monomeric member of the small leucine-rich repeat protein (SLRP) family ^42^, as significantly upregulated in patients who developed bone metastases compared to those who developed metastases at other metastatic sites or who did not develop metastases. The transcript of OMD was previously identified as enriched in breast cancer bone metastases compared to breast cancer metastases in other organs ^10^ and in prostate cancer bone metastases ^19^. These findings suggest that OMD may be a pan-cancer biomarker for bone metastases. OMD regulates important cancer pathways, including Ras, P53, Jun/Fos, MYC, and NFκB signalling ^24^ but little is known about its role in cancer phenotypes. Although our data exclude a primary role for OMD in the tested functions of fibroblasts and osteoclasts using available cell models, OMD may play a role in promoting bone turnover, which is an essential step in osteosclerotic bone metastases ^25^. OMD is known to reduce apoptosis in osteoblasts ^43^, thus supporting a role for OMD in promoting new bone formation and cell survival. Indeed, in our patient cohort upregulated OMD expression correlates with upregulation of serine 118 phosphorylation on pro-apoptotic protein BCL2-associated agonist of cell death (BAD) (Table S5). This site prevents BAD interaction with BCL2 Like 1, thus preventing apoptosis ^44^. We establish here that OMD regulates cell migration in both ER+ and ER-cell lines and that secretion of OMD is from CAFs, not tumour cells. We validate this by showing that CAFs secrete biologically relevant concentrations of OMD^45^, which stimulate tumour cell migration. In support of this mechanism, recent data demonstrate cancer-associated fibroblasts are strongly associated with bone metastasis in over 300 early invasive ductal breast cancer patients ^46^.

Secreted OMD ^24^ engages cell surface receptors, such as the BMP2 receptor complex ^45^, and integrin α_V_β_3_, both of which are associated with bone metastasis ^47,48^. In support of this idea, we identified BMP2-induced kinase (BMP2K) and FAK activity downstream of OMD (Table S11 and data not shown). Crosslinking^49^ and cell-surface capture mass spectrometry^50^, or functional kinase profiling assays could be used to identify other candidates and elucidate the signalling pathways downstream of OMD in breast and other cancer cells. Phosphoproteomics analysis identified a correlation between elevated OMD expression and increased activation of the kinase CDK1 in both cell models and patient samples. Therefore, OMD expression may inform clinicians as to which patients could benefit from CDK1 inhibition. Indeed, we demonstrated the independent prognostic impact of a OMD/CDK1 signature on metastatic recurrence in over 2,600 breast cancer patients, with the strongest predictive value observed in ER+ cases. This would require further validation at the protein level to develop an effective biomarker to select for OMD/CDK1 therapy to prevent bone metastasis.

Osteoblasts express and secrete OMD^51,52^, and it is therefore feasible that when cancer cells arrive in the bone microenvironment, osteoblast-secreted OMD enables cancer cell colonisation. Furthermore, CDK1 phosphorylates and activates RUNX2, which can induce a variety of bone-related genes ^53,54^, and we speculate that OMD may drive bone metastasis through CDK1-RUNX2 ^55,56^.

We validated that the OMD-CDK1 axis is a driver of cell viability and motility, consistent with the role of CDK1 not only in cell cycle regulation, but also in actin remodelling, a key process in cell migration ^57^. We also found increased phosphorylation of vimentin on Serine 56 (Table S5), which regulates cancer cell invasion ^58^, in patients who developed metastases compared to those who did not (Table S5). The intriguing role of CDK1 as a driver of cell migration and potential metastasis deserves further investigation in breast and other cancers. Indeed, CDK1 mRNA is upregulated not only in several transcriptomics datasets of breast cancer samples compared to normal breast tissue samples ^59^, but also in melanoma ^60^, thyroid cancer ^61^ and acute myeloid leukaemia ^20^. Additionally, our study investigating tumour cells derived from patients with bone metastases showed a relationship between OMD protein expression, CDK1 activation (reduced Y15 phosphorylation) and reduced cell viability in response to CDK1 inhibitor. As discussed above, OMD may activate BAD to drive the survival of breast cancer patient cells during bone metastasis. Finally, we demonstrate higher bone colonisation after intracardiac injection of OMD-overexpressing tumour cells and that this can be abrogated by treatment with a CDK1 inhibitor. Overall, our data suggest that CDK inhibition could serve as a novel therapeutic route to decrease motility, viability and bone metastasis in breast cancer. However, the CDK1 inhibitor RO-3306 has been used in few studies *in vivo* ^62,63^ and no CDK1-selective inhibitor has been tested in clinical trials in breast or other cancers ^64^. Future studies should aim at testing other CDK1-selective inhibitors or CDK1 inhibition in combination with other signalling inhibitors. For instance, among the phosphorylated sites significantly upregulated in patients who developed metastases compared to those who did not, we also found phosphorylated ser 641 on catenin alpha 1 (CTNNA1) (Table S5) which activates beta catenin signalling resulting in tumour cell invasion ^65^ and phosphorylated Thr185/Tyr187 on MAPK1 which are hyperactive in metastatic TNBC ^13^ (Table S5). Therefore, our data provides the rationale for the targeting of MAPK and beta catenin signalling pathways in combination with or independently of CDK1 inhibition to prevent metastasis.

In conclusion, the identification of the OMD-CDK1 axis from patient-derived samples with complete clinical information suggests that similar studies can be performed in any cancer setting including across multiple cancers to identify novel biomarkers, organotropic metastatic drivers, and therapeutic targets. Here, OMD was discovered to be significantly upregulated in primary breast cancer biopsies of patients who developed bone metastases within five years of diagnosis, compared to those who were distant disease free at 5 years. OMD significantly increased cell migration and cell viability as well as bone colonisation of breast cancer cells and of breast cancer cells derived from patients with bone metastases. Migration, viability and bone colonisation induced by CAF-derived OMD were CDK1-dependent and thus highlight a key tumour-stroma interaction in bone metastasis establishing the OMD–CDK1 axis as a potential therapeutic target in metastatic breast cancer.

## Resource Availability

### Data and code availability

All raw MS data were generated by the authors and deposited to the ProteomeXchange Consortium, via the PRIDE partner repository with the dataset identifier PXD041719 (Username; Password: 4PS3MTqM); PXD041720 (Username; Password: Tr6QtMFt); PXD042764 (Username; Password: rsDlM5NX).

The R scripts related to this publication are freely accessible at https://github.com/Joseph-Parsons/OMD/.

## Acknowledgments

Funding: This work was supported by Engineering and Physical Sciences Research Council grant EP/S022201/1 (SC), CR-UK Non-Clinical Training Award A27445 (JP), Medical Research Council grant MR/T016043/1 (PF, DB, JF, MPS, JW), Breast Cancer Now (HH, KS, PO, DL), Stanley Center for Psychiatric Research at Broad Institute (KT), NIHR Manchester Biomedical Research Centre (TK, NA), Cancer Research UK Manchester Institute Histology Core Facility (CB), Yorkshire Cancer Research, Western Park Cancer Charity (PO), National Institute for Health Research (COB), Cancer Research UK, National Institute for Health Research, UK Research and Innovation, Breast Cancer Now (RBC), Wellcome Trust 107636/ Z/15/Z and 107636/Z/15/A, the Biotechnology and Biological Sciences Research Council BB/X001970/1 and Medical Research Council MR/T016043/1, the NNF Young Investigator Award call 2022, NNF22OC0070845 (CF). For the purpose of open access, the author has applied a CC BY public copyright licence to any Author Accepted Manuscript version arising from this submission.

We thank the CRUK Manchester Institute Histology and Visualisation, Imaging and Analysis facilities for their support with immunohistochemical optimisation and visualisation, the University of Manchester Bio-MS, Genome Editing, and Imaging facilities for their continuing support throughout the project. We thank Dr Hurlstone, University of Manchester for providing invaluable reagents and protocols.

## Author contributions

S.C., J.P. performed the research and interpreted data. H.H., T.K., P.F., K.S., D.B., J.F., J.F., C.B., N.A., M.P.S, P.O., D.L., J.B. contributed to the research and interpreted data. J.W., K.T., M.J.D., B.G. supported and contributed to the bioinformatic analyses. C.O.B. devised and supervised the project and collated information on patient samples for mass spectrometry-based analysis. CF devised and supervised the project, designed mass spectrometry and *in vitro* experiments, and obtained funding for the project. RBC devised and supervised the project, designed *in vitro* and *in vivo* experiments, and obtained funding for the project. All authors reviewed and approved the manuscript before submission.

## STAR Methods

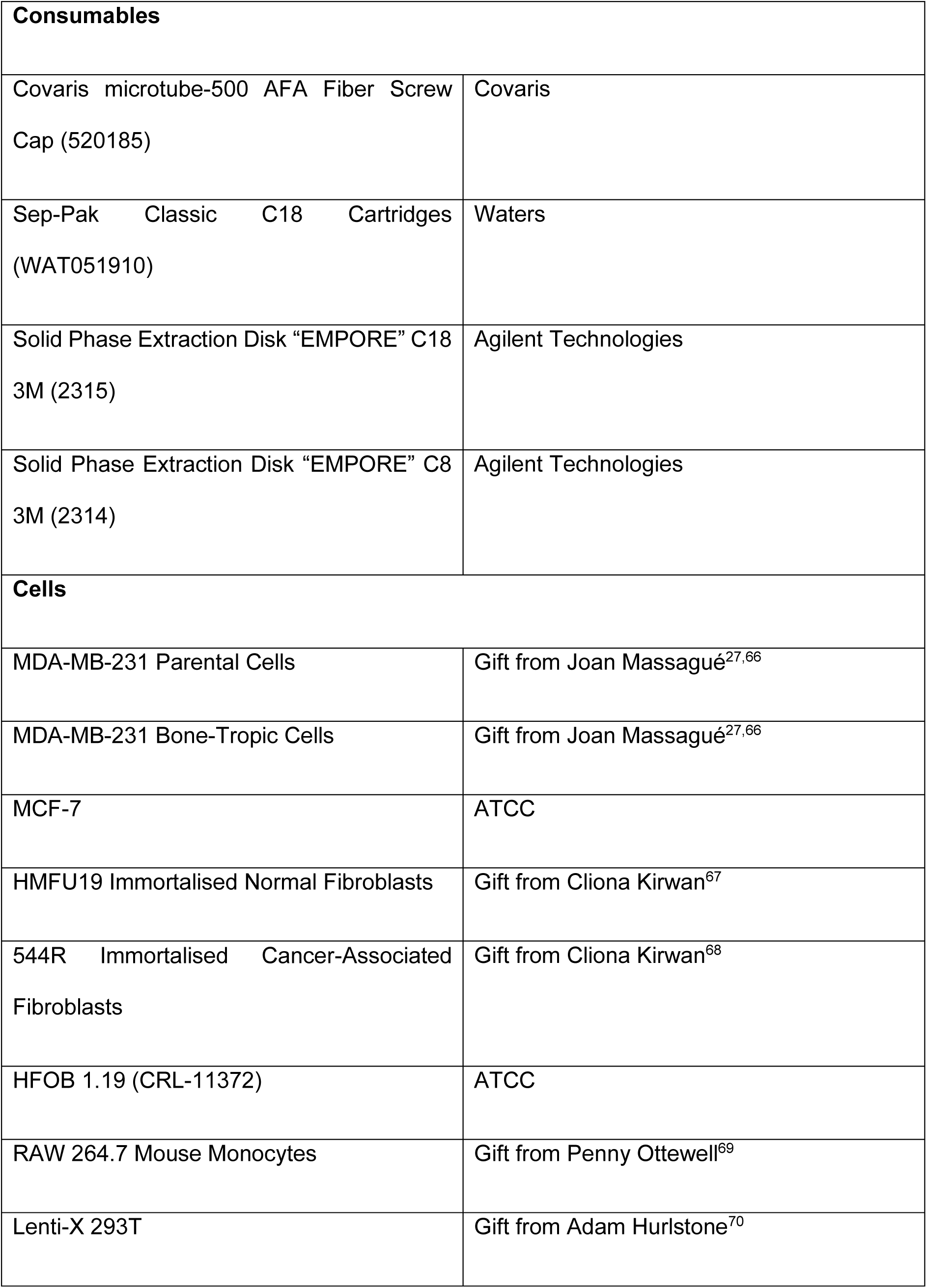
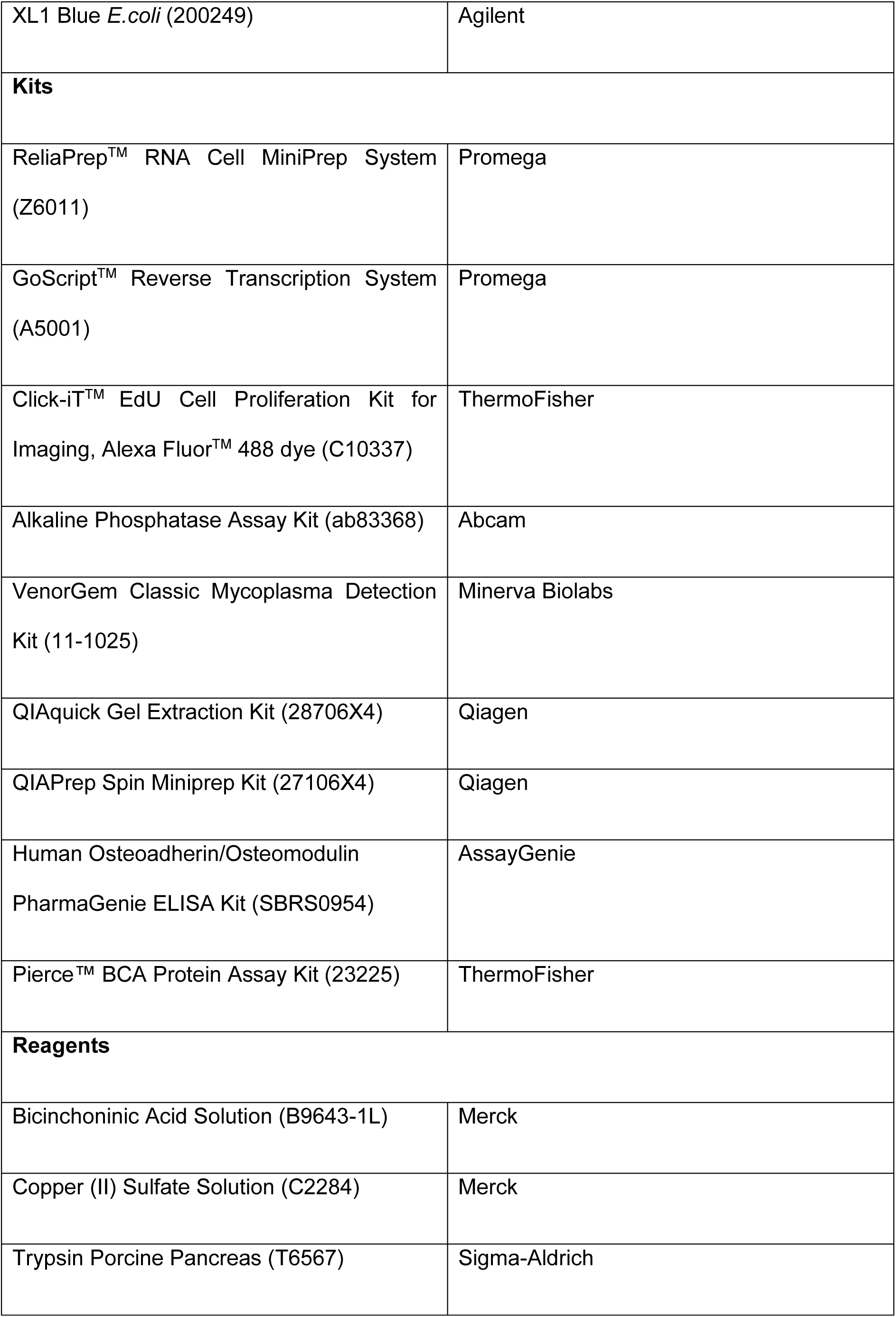
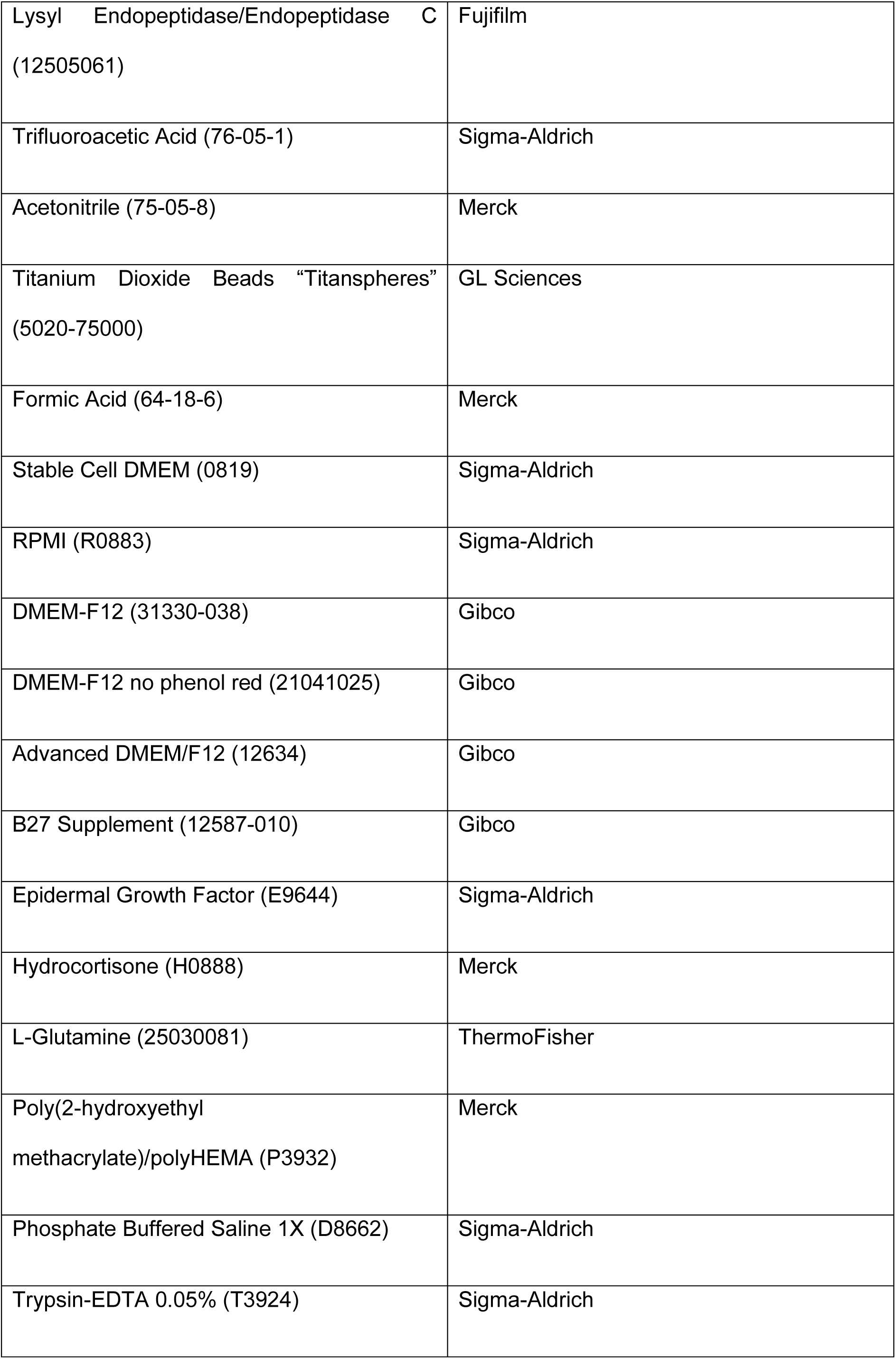
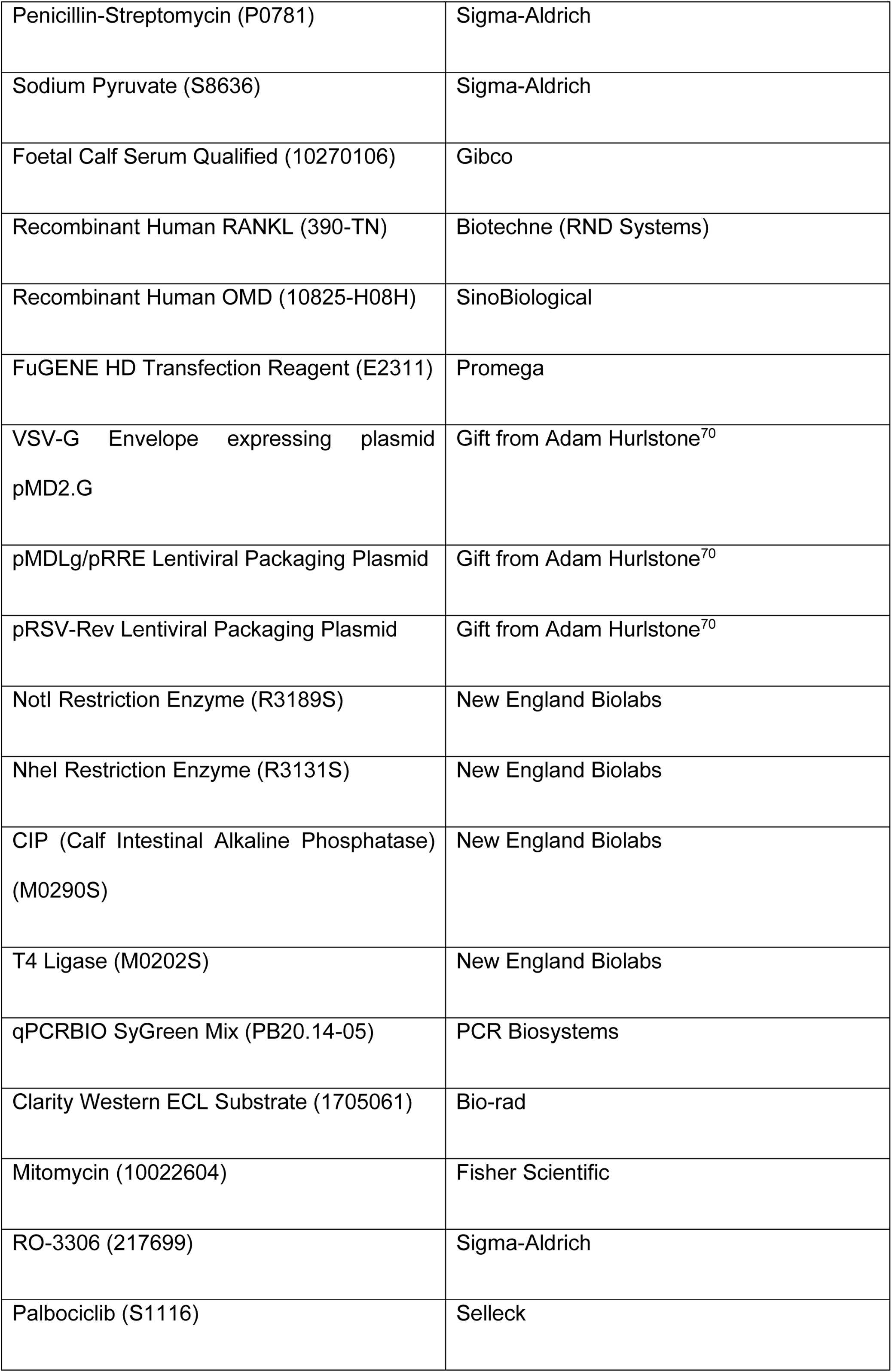
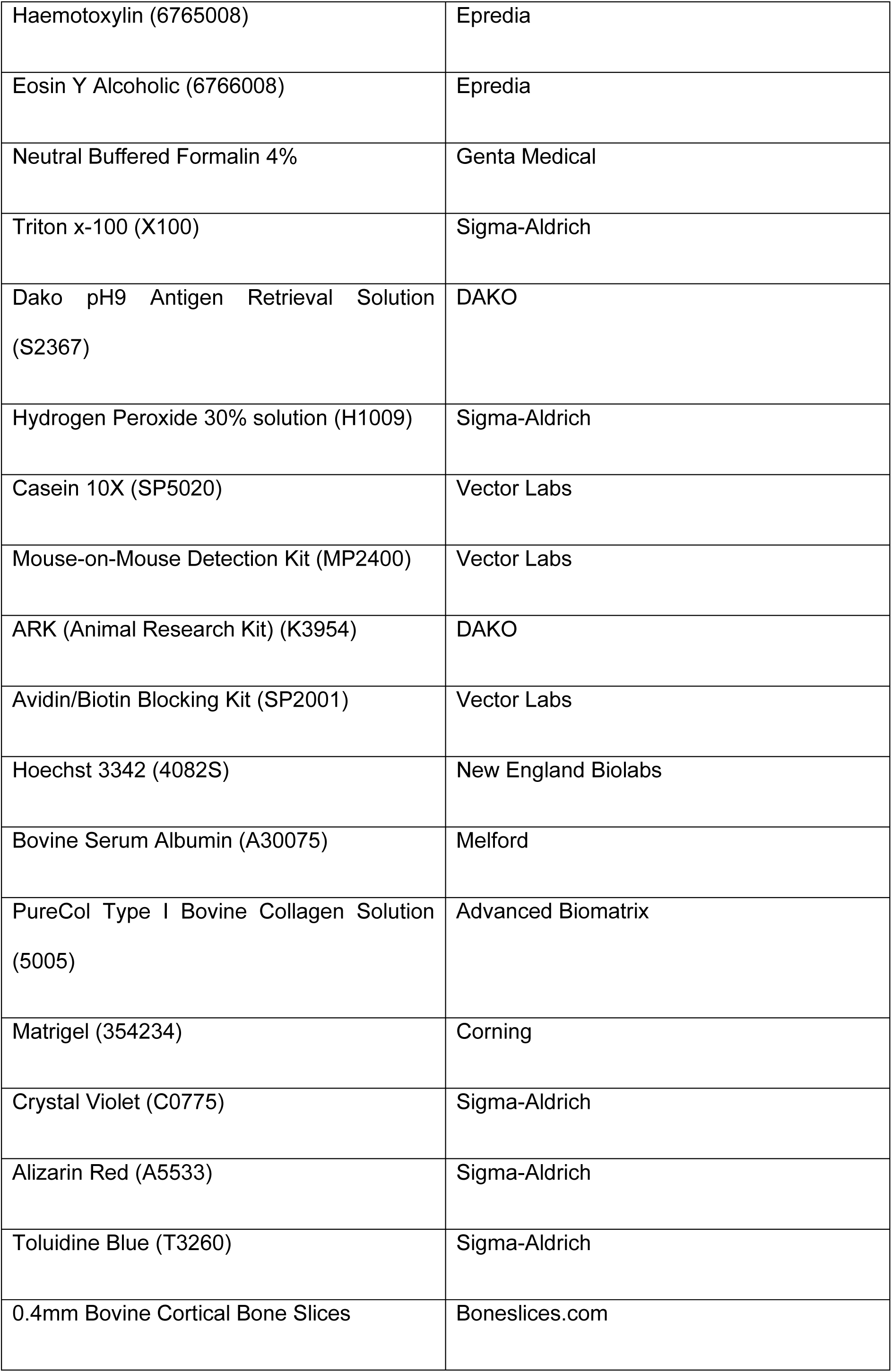
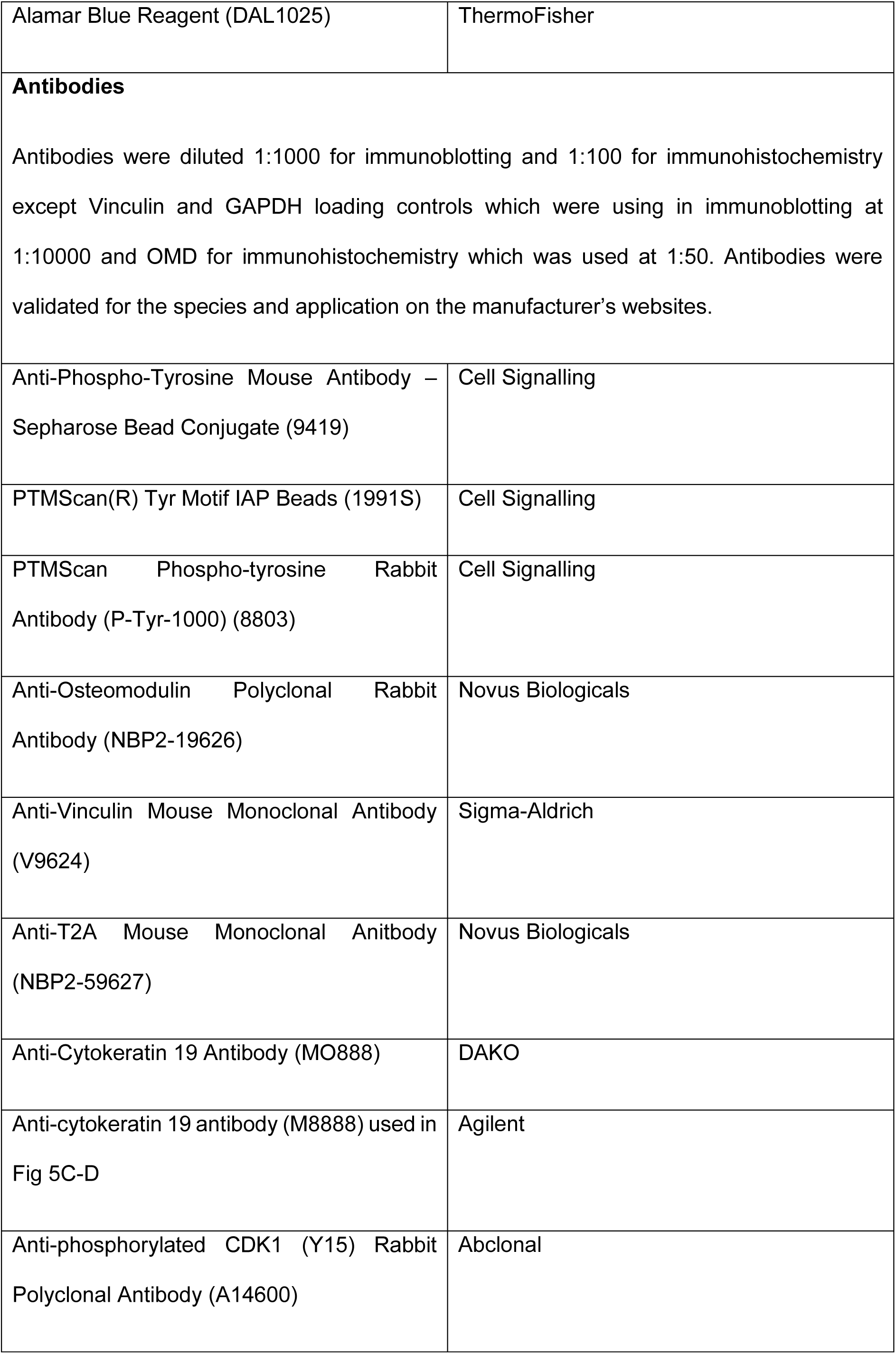
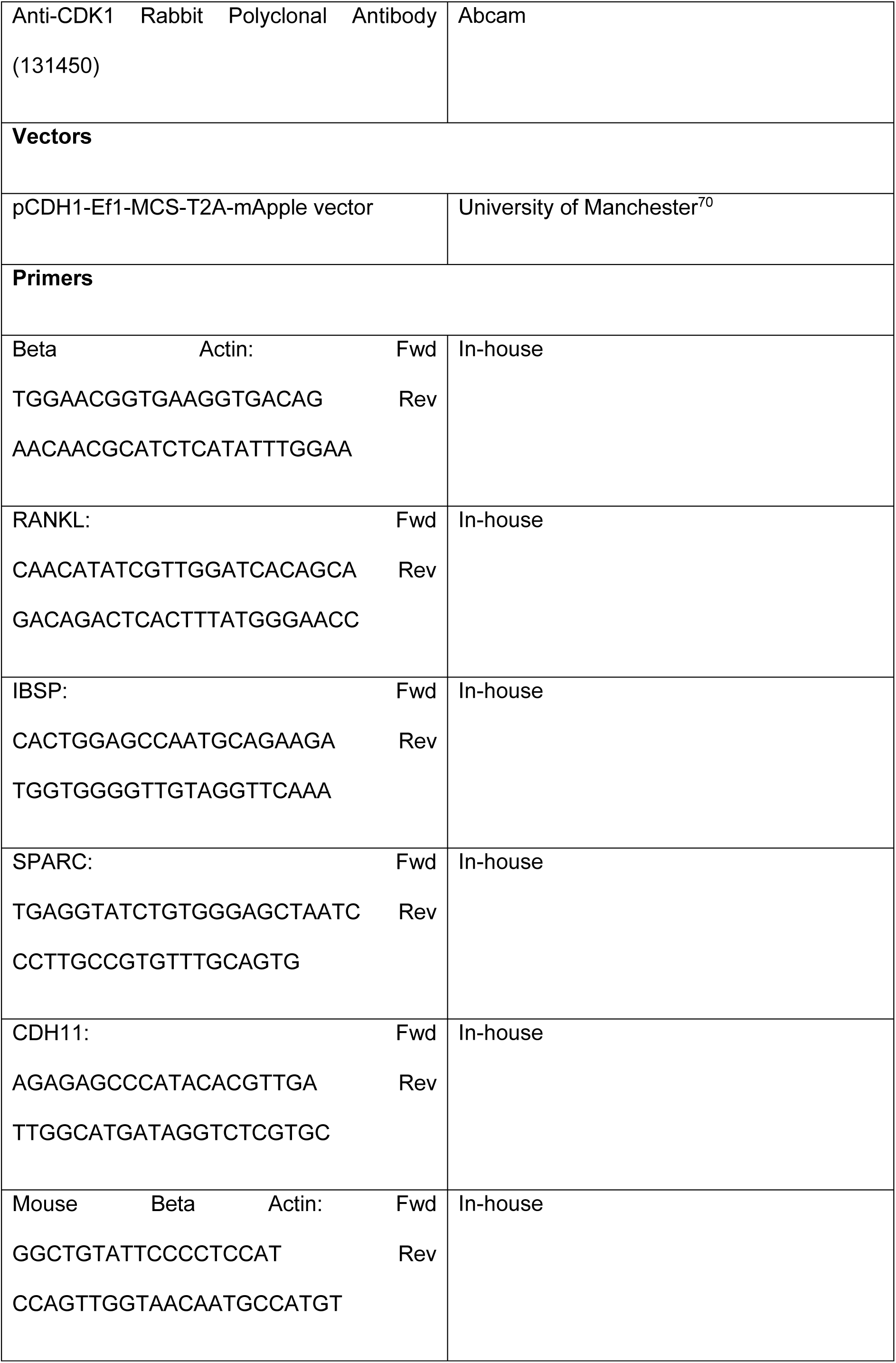
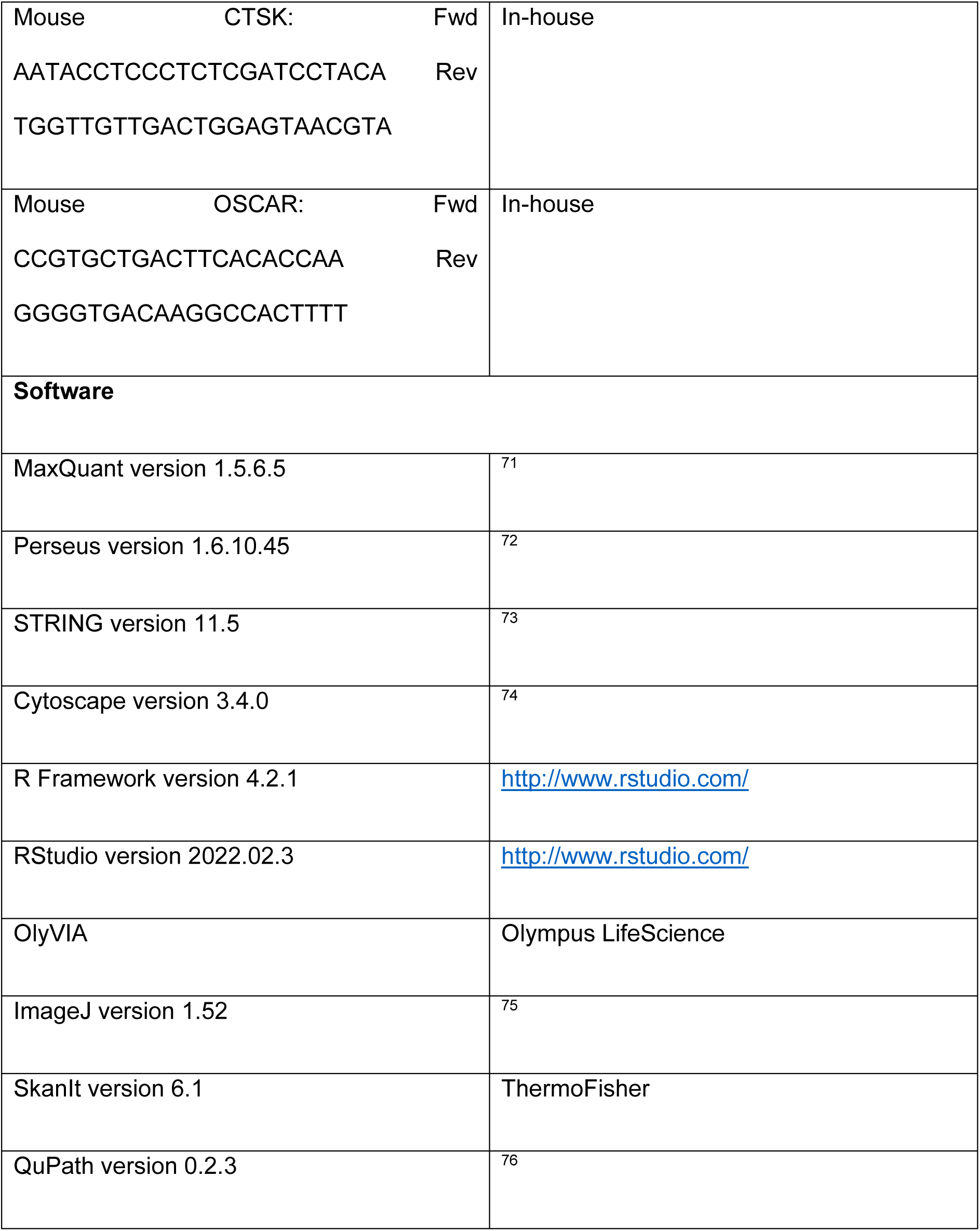
Key Resources Table.

### Ethics approval

Early breast cancer samples were collected from patients undergoing surgical resection of the primary tumour at the Manchester University NHS Foundation Trust through the Manchester Cancer Research Centre Biobank. Metastatic fluids (ascites or pleural effusions) drained from advanced breast cancer patients were collected at the Christie and South Manchester Hospitals NHS Foundation Trusts through the Manchester Cancer Research Centre Biobank. Fully informed consent from all patients was obtained in accordance with local National Research Ethics Service guidelines (study numbers: 05/ Q1402/25, 05/Q1403/159, and 12/ROCL/01). Patient samples from the University of Sheffield sponsored AZURE trial were obtained with fully informed patient consent. AZURE was approved by the Clinical Trials Committee; controlled trials number ISRCTN79831382^29^.

### *In vivo* Experiments

All *in vivo* experiments were performed in accordance with the Animals (Scientific Procedures) Act 1986 under the UK Home Office project licence PCFEAC0D4 and study protocols were approved by the CRUK Manchester Institute Animal Welfare and Ethical Research Board (AWERB). 100,000 PAM, PAO, or BOM cells in 50µl PBS were injected into the left cardiac ventricle of 4 athymic nude mice (sourced from Envigo) per condition using a 28-gauge needle. Mice were monitored for 25 days before culling. For CDK1 inhibition studies, 4 days after intracardiac injection of 100,000 PAO cells, 4 athymic nude mice were dosed with 225µl of RO-3306 solution (final concentration 4mg/kg, SeleckChem) dissolved in 40% PEG300-1% Tween80-3% DMSO in ddH_2_O given 3 times per week by oral gavage and compared to 4 vehicle control mice. Mice were culled 25 days after intracardiac injection of breast cancer cells. Lungs (only for the first study) and one femur were formalin-fixed for immunohistochemical analysis, and one femur was collected in DMEM with FCS, 10% Pen-Strep and 1mM Sodium pyruvate and bones were flushed. To flush bones, the ends of mouse femurs were clipped using scissors and ground down and the bone marrow was flushed using 5mL complete DMEM using a 25-gauge needle. The ground bone ends and bone marrow were filtered through a 100µm filter which was washed with 5mL complete DMEM. Filtered bone cells were pelleted by centrifugation at 600xg at 4°C before filtering through a 40µm filter.

### Sample Preparation for Proteomics and Phosphoproteomics

Fresh frozen primary breast cancer patient samples were ground using a pestle and a mortar filled with liquid nitrogen to prevent tissue thawing. Technical replicates of 9 samples were used to assess reproducibility of the method. For both fresh frozen primary breast cancer patient samples and metastatic effusion samples, 6M guanidine hydrochloride in 100mM Tris pH 8.5 at 99°C was immediately added to the ground tissue and samples were heated at 99 degrees for 10 minutes in an Eppendorf shaker at 600rpm. Lysates were transferred to Covaris sonicator tubes and sonicated in a Covaris E220 (Covaris) three times for three minutes cycles with peak power of 500, duty factor 30%, 200 cycles/burst and 150 average power. Protein concentration was assessed using bicinchoninic acid assay (ThermoFisher). 750ug of protein was digested to peptides using Endopeptidase C for 1 hour before dilution with 25mM Tris pH 8.5 and further digestion with trypsin overnight. Trifluoroacetic acid (TFA) was added to 1% concentration to eliminate further trypsin activity. Samples were desalted using SepPak columns (Waters), eluted using 50% acetonitrile (ACN) dried using an Eppendorf Concentrator, resuspended in 10mM ammonium hydroxide (NH_4_OH) pH 10 and fractionated using a StageTip-based reverse phase fractionation method. StageTips were initially activated with methanol, 5mM NH_4_OH 50% ACN and 10mM NH_4_OH before loading samples, taking a flow-through fraction, and subsequently eluting with 1% ACN and 7% ACN into different fractions. A 50% ACN fraction is taken to remove any remaining peptides bound to the C18 disks. The same fractionation with a flow-through, 1%, 7% and 50% ACN fraction was performed on a second tip. The flow through and 50% fractions were combined, both 1% fractions were combined and both 7% fractions were combined. 5% of each sample was retained for proteome analysis. The remaining 95% was acidified with 6% TFA in 50% ACN before incubation with titanium dioxide coated beads (GL Sciences) in 20mg/ml dihydroxybenzoic acid (DHB) resuspended in 80% ACN, 6% TFA and deionised water, as previously described^17^. Beads were centrifuged onto a C8 disc (EMPORE) and then were washed with 10% ACN 6% TFA followed by 40% ACN 6% TFA and 80% ACN 6% TFA before subsequently being eluted using 5% ammonia and 10% ammonia in 25% ACN. Both proteome and phosphoproteome samples were loaded onto C18 StageTips (EMPORE) for an additional desalting step before elution using 40% ACN. Samples were then reduced to 5ul using an Eppendorf Concentrator and resuspended in 25ul 0.1% Formic Acid, 5% TFA before analysis on the mass spectrometer.

MDA-MB-231 cells (three independent biological replicates of 231-mA, 231-OMD, 231-Bo and 231-Bo OMD) were lysed with 6M GuHCl in 100mM Tris pH 8.5 heated to 99°C. Lysates were sonicated and protein concentration assessed as above. 1.5mg of protein per sample was digested and desalted as above. Dried samples were resuspended in 50mM MOPS pH 7.2, 10mM Sodium Phosphate, 50mM Sodium Chloride (NaCl) and dissolved overnight. Samples were clarified by centrifugation and clarified peptides were incubated in a new tube with immobilised phosphorylated tyrosine antibody beads (Cell Signalling) and incubated for 2 hours at 4°C. Beads were washed five times with the MOPS buffer and twice with 50mM NaCl. Phosphorylated tyrosine-enriched peptides were eluted from the beads three times using 0.1% TFA. Unbound samples were processed as above with 5% of sample retained for proteome analysis and the rest being used for phosphorylated peptide enrichment. Proteome, phosphoproteome and phosphorylated tyrosine-enriched samples were loaded onto C18 StageTips, eluted using 40% ACN, reduced to 5ul and resuspended in 0.1% Formic acid, 5% TFA before analysis on the mass spectrometer.

Cells from metastatic effusions were pelleted by centrifugation, the supernatant was removed, and cells were lysed and processed for proteomic analysis via the same processes as MDAMB231 cell lines, described above.

### Mass Spectrometry Analysis

Mass spectrometry analysis was performed as described^17^. Briefly, purified peptides were analysed by LC-MS/MS using an UltiMate 3000 Rapid Separation LC (ThermoFisher) coupled to an Orbitrap Exploris 480 mass spectrometer (ThermoFisher). Mobile phase A was 0.1% FA in water, and mobile phase B was 0.1% FA in ACN and the column was a 75 mm x 250 μm inner diameter 1.7 μM CSH C18, analytical column (Waters). A 1 μl aliquot of the sample (for proteome analysis) or a 3 μl aliquot was transferred to a 5 μl loop and loaded on to the column at a flow of 300 nl/min at 5% B for 5 and 13 min, respectively. The loop was then taken out of line and the flow was reduced from 300 nl/min to 200nl/min in 1 min., and to 7% B. Peptides were separated using a gradient that went from 7% to 18% B in 64 min., then from 18% to 27% B in 8 min. and finally from 27% B to 60% B in 1 min. The column was washed at 60% B for 3 min. and then re-equilibrated for a further 6.5 min. At 85 min, the flow was increased to 300nl/min until the end of the run at 90min. Mass spectrometry data were acquired in a data-dependent manner for 90 min in positive mode. Peptides were selected for fragmentation automatically by data-dependent analysis on a basis of the top 8 (phosphoproteome analysis) or top 12 (proteome analysis) with m/z between 300 and 1750Th and a charge state of 2, 3 or 4 with a dynamic exclusion set at 15 s. The MS resolution was set at 120,000 with an AGC target of 3e6 and a maximum fill time set at 20ms. The MS2 resolution was set to 60,000, with an AGC target of 2e5, and a maximum fill time of 110 ms for Top12 methods, and 30,000, with an AGC target of 2e5, and a maximum fill time of 45 ms for Top8 analysis. The isolation window was of 1.3Th and the collision energy was set at 28.

### Data Analysis

Raw files were analysed by the MaxQuant software suite^71^ (version 1.5.6.5) using the integrated Andromeda search engine. Proteins were identified by searching the HCD-MS/MS peak lists against a target/decoy version of the human UniProt Knowledgebase database that consisted of the complete proteome sets and isoforms (v.2019; https://uniprot.org/proteomes/UP000005640_9606) supplemented with commonly observed contaminants such as porcine trypsin and bovine serum proteins. Tandem mass spectra were initially matched with a mass tolerance of 7 ppm on precursor masses and 0.02 Da or 20 ppm for fragment ions. Cysteine carbamidomethylation was searched as a fixed modification. Protein N-acetylation, N-pyro-glutamine, oxidized methionine and phosphorylation of serine, threonine and tyrosine were searched as variable modifications. Protein N-acetylation, oxidized methionine and deamidation of asparagine and glutamine were searched as variable modifications for the proteome experiments. Label-free parameters were used. False discovery rate was set to 0.01 for peptides, proteins, and modification sites. Minimal peptide length was six amino acids. Site localization probabilities were calculated by MaxQuant using the PTM scoring algorithm. The datasets were filtered by posterior error probability to achieve a false discovery rate below 1% for peptides, proteins, and modification sites. Only peptides with Andromeda score > 40 were included in the downstream analysis.

For patient sample analysis, proteome quality control included proteins with more than 2 razor+unique peptides and greater than 5% protein sequence coverage. For the phosphoproteome, localisation probability of 0.7 or higher was required for each site. Samples with fewer than 2000 proteins or 2000 phosphorylated sites were excluded. Only proteins and sites that were present in at least 80% of either non-metastatic or metastatic samples were included in the final analysis. For cell line and metastatic effusion samples, we used the same criteria as above and we included only proteins or phosphorylated sites that were present in all samples.

All statistical and bioinformatics analysis was done using freely available software and packages: Rstudio (R version 4.2.1, Rstudio version 2022.02.3) (http://www.rstudio.com), Perseus (version 1.6.10.45)^72^, STRING (version 11.5)^73^, and Cytoscape (version 3.4.0)^74^. Differential expression analysis performed separately on proteome and phosphoproteome using Rstudio package DEP (Bioconductor version 3.15). All peptide intensities were normalised using the normalizeQuantiles function in the R package Limma^77^. Normalised peptides were subjected to left-centred imputation and differential expression analysis using the R packages DEP (PMID: 29446774) and Limma^77^. Differentially expressed proteins or phosphorylated sites were defined in patient samples as those with a p value of less than 0.05 and a Log2 fold change of less than –1.5 or greater than +1.5, and in cell lines as those with a p value of less than 0.05 and a Log2 fold change of less than –1 or greater than +1. Hierarchical clustering and Vulcano plots analyses were performed using Perseus. Protein-protein interactions networks were generated using STRING. Data were visualised with Cytoscape.

### Analysis of the GEO datasets and single cell RNA seq dataset

The analysis of OMD expression in GEO datasets shown in Fig. 1F was performed using Table 2 from reference^10^. Single cell RNA seq dataset was analysed using the online platform CELLxGENE^28^.

### Generation of a Database of Breast Cancer Metastasis-Associated Genes and Proteins

This database was generated using the PubMed search term “Breast Cancer Metastasis” and collecting the first 300 most relevant texts. Genes and proteins dysregulated in breast cancer metastasis were extracted from papers and presented in the database with gene name, protein name(s), a short summary of their contribution to breast cancer metastasis, the reference for the energy, any specific organotropism associated with the entry and the level at which the dysregulation occurs e.g., genomics, transcriptomic or proteomic.

### Kaplan-Meier analysis

Gene IDs for OMD (205907_s_at, 205908_s_at) and CDK1 (203214_x_at, 210559_s_at, 203422_at, 203213_at) were used as input to the breast cancer mRNA gene chip dataset at kmplot.com^34^ to evaluate the effects of the two-gene signature on distant metastasis-free survival. All patients and patients with estrogen receptor positive subtype were dichotomized according to the auto selected best cutoff percentile of the mean expression of the gene signature and the threshold set at 180 months.

### Multivariate Survival Analysis

To assess the prognostic significance of a two-gene signature in breast cancer, we performed a comprehensive search in the GEO (https://www.ncbi.nlm.nih.gov/geo/) and EGA (https://ega-archive.org/) repositories. Transcriptomic datasets with DMFS data, with a minimum of 30 samples, were included and focused on those generated using the platforms GPL96, GPL570, and GPL571. The datasets used in this analysis were GSE2990, GSE3494, GSE6532, GSE7390, GSE11121, GSE5327, GSE9195, GSE16446, GSE17907, GSE19615, GSE20685, GSE26971, GSE16716, E-TABM-43, GSE45255, GSE25066, GSE22093, GSE65194, GSE69031, GSE58812, and GSE61304. This selection ensured the use of a consistent set of 22,277 genes across datasets, allowing for uniform sensitivity, specificity, and dynamic range. Each dataset underwent rigorous normalization and quality control. Initially, normalization was conducted using MAS5, a method chosen for its superior performance in previous comparisons with RT-PCR-validated expression data. MAS5 also enables the normalization of individual samples independently, which prevents the recalibration of values when adding or removing samples. To reduce batch effects, a second normalization step was applied, scaling the mean expression of overlapping probes to a value of 1,000 across all arrays. Redundant samples with identical gene expression profiles were identified and excluded, retaining only the earliest version to ensure unique biological representation. Quality control metrics were assessed across five parameters, including background signal, raw Q values, the percentage of present calls, bioBCD spike-in controls, and the GAPDH/ACTB 3-to-5 ratio. Samples were deemed acceptable if they met the expected range for all parameters, while outliers and biased arrays, failing two or more criteria, were excluded from downstream analyses. Following this process, 2,828 unique samples with high-quality data were retained for subsequent analysis. The two-gene signature was constructed by calculating the mean expression of the probes corresponding to OMD (osteoadherin) and CDK1 (cyclin-dependent kinase 1). The signature utilized six probe sets: 205908_s_at, 205907_s_at (OMD), and 203214_x_at, 210559_s_at, 203422_at, 203213_at (CDK1). Survival analysis was performed to evaluate the association between the signature and distant metastasis-free survival (DMFS). The mean expression of the signature was assessed across potential cutoff values spanning the interquartile range, with statistical significance determined using Cox proportional hazards regression models. False Discovery Rate (FDR) correction was applied using the Benjamini-Hochberg method. Subgroup analyses were conducted for estrogen receptor-positive (ER+), triple-negative breast cancer (TNBC), and all tumors collectively. The independent prognostic value of the signature was further assessed using multivariate Cox regression, accounting for available clinical and pathological parameters including HER2 expression, lymph node status, tumor grade, tumor size, and patient age.

### Immunohistochemistry

500,000 bone marrow cells were pelleted onto glass slides using a Shandon Cytospin 3 centrifuge at 600rpm. Cytospin samples were fixed in 4% neutral-buffered formalin in PBS and permeabilised with 0.25% Triton X-100. Endogenous peroxidases were blocked with hydrogen peroxide (10%), samples were blocked with Casein. Samples were incubated with 200µg/ml anti-Cytokeratin 19 primary antibody (CK19) was biotinylated using DAKO Animal Research kit (ARK) and incubated on samples for 30 minutes at room temperature. ARK kit anti-biotin secondary antibodies were incubated on samples for 30 minutes before staining with DAB stain to produce a colorimetric response. Gills I haemotoxylin was used for counterstaining and samples were coverslipped before imaging. TBST washes were performed between each step. Whole slide imaging was performed using Olympus VS200 MTL (Olympus Life Science) in conjunction with an Olympus UPLXAPO20X objective lens and 20x magnification. Lung metastatic tumour lesions, identified in collaboration with Visualisation, Imaging and Analysis Core Facility at Cancer Research UK Manchester Institute, of greater than 100µm in diameter (in one orientation) were counted using OlyVIA to assess the number of macrometastases in the lungs. For quantification of cytokeratin 19-positive cells in cytospins of the bone marrow flushes for figure 6C and D, slides were scanned and images were opened in OlyVIA and sample coordinates were determined and a random number generator was used to select coordinates in which five fields of view at 10X magnification were used to count the number of CK19-positive cells. For quantification of cytokeratin 19-positive cells in cytospins of the bone marrow flushes for figure 6E and 6F, slides were scanned, and images were opened in OlyVIA and all 4 quadrants of the cytospins were scored to enumerate the CK-positive cells in control and CDK1 inhibitor treated mice. Immunohistochemical staining of patient samples for osteomodulin required sample dewaxing in xylene, washing sequentially with 100% industrial denatured alcohol (IDA), 90% IDA and 70% IDA and deionised water. Antigen retrieval was performed in DAKO pH9 antigen retrieval solution using an EZ Retriever system (BioGenex) at 98°C for 15 minutes. Samples were washed with deionised water and PBS. Endogenous peroxidases were blocked by incubation with 10% hydrogen peroxide in PBS for 10 minutes. Slides were incubated with Casein for 1 hour at 37°C. Anti-OMD primary antibody was added at 1:50 for 1 hour at 37°C. Slides were washed three times with PBS. Envision anti-mouse secondary fluorescent antibody was added to slides for 30 minutes at 37°C. Slides were washed twice with PBS. DAB was added (1 drop DAB in 1ml diluent) to slides for 5 minutes before washing with deionised water and staining for 10 seconds with Gills I haemotoxylin. Slides were washed with water, 70% IDA, 90% IDA and 100% IDA. Slides were air-dried and scanned as above. Analysis for our cohort of patient samples was performed by consultant pathologist Nisha Ali who selected five fields of view per slide and assessed the cell types that were positively stained within each field of view based on morphology. Haemotoxylin and Eosin staining of breast cancer patient samples was performed by Cancer Research UK Manchester Institute Histology facility and epithelial percentage in each section was calculated by pathologist Nisha Ali who identified epithelial cells by morphology and estimate the area covered by epithelial cells compared to the total size of tissue on the slide. Analysis of primary tumours from patients recruited into the trial ^29^ were performed on TMAs constructed for the purpose of this study. Identification of positively stained areas was conducted using QuPath version 0.2.3 using the positive pixel count module.

### Quantitative RT-PCR

RNA was isolated from cells at 70-90% confluency using a ReliaPrep^TM^ RNA Cell Miniprep system according to manufacturer’s instructions. Isolated RNA was reverse transcribed using a GoScript^TM^ Reverse Transcription System according to manufacturer’s instructions. Specific genes within complementary DNA (cDNA) were amplified using gene-specific primers and a 2X SYBR Green mixture. The qPCR machines used were the Agilent Mx3000P and the Aria Mx Real-time qPCR System. Relative expression was quantified using the Delta-Delta-CT method with β-actin as a housekeeping gene.

### Cell Lysis and Immunoblotting

Cell lines were lysed with SDS sample buffer supplemented with beta mercaptoethanol and samples were heated at 95 degrees for 30 minutes. Patient samples were pelleted and lysed at 4°C on a rotating wheel for 1 hr. Lysates were centrifuged at 12,000g to remove debris. Supernatant was quantified using Pierce™ BCA Protein Assay (ThermoFisher) following the manufacturers protocol. Lysates were normalised to the same concentration and boiled at 95°C for 5 minutes. All proteins were resolved by SDS-PAGE and transferred onto nitrocellulose membranes. Specific antibodies were used to probe for proteins of interest and peroxidase-labelled secondary antibodies recognised primary antibodies. An enhanced chemiluminescence kit (Biorad) was used to assess expression and blots were visualised using a Universal Hood II Gel Molecular Imaging System (Biorad).

Densitometric analysis was performed using ImageJ (version 1.52). Tiff images exported from ImageLab (Biorad) were converted to 8bit images. The square selection tool was used to highlight the first lane and pressing Ctrl+1 allows the dimensions of this box to be moved over each lane so that the selection size was identical. Background was subtracted from the generated base of the intensity curves using the line tool. The magic wand tool was used to highlight the area underneath each peak providing a quantification of the area under the intensity curve. Intensity values were divided by the loading control from the same lane to normalise peak intensities.

### EdU Incorporation Assay

Cells were incubated with 20µM 5-ethynyl-2’-deoxyuridine (EdU) for 4 hours at 37°C. Cells were fixed with 4% paraformaldehyde (PFA) in PBS and processed according to manufacturer’s instructions. After staining, cells were counterstained with 5ng/ml Hoechst 3342 for 30 minutes and counted using a Leica microscope system. Approximately 100 stained nuclei were counted before changing to a FITC filter and counting the number of cells that also stained for Alexa488.

### Collagen Contractility Assay

Flat-bottomed 96 well plates were coated with 100µl of sterile-filtered 3% bovine serum albumin (BSA) in PBS for at least 1 hour at 37°C before aspiration. 50,000 HFOB 1.19 cells or 100,000 of MDA-MB-231, HMFU19 or 544R CAFs were resuspended in 10ul of their respective medias. 7mg/ml Bovine type I collagen solution with 10% FCS at pH 7.7 was made. 90µl of the collagen mixture was mixed with 10µl cell suspension and added to a coated well and allowed to set at 37°C for 1 hour. 100µl of media was added. Any drugs or recombinant protein used were added in this 100µl such as recombinant OMD on top of the collagen plug. Samples were incubated at 37°C for 72 hours before imaging using a GelCount image system. Images were analysed using area measurements on ImageJ.

### Boyden Chamber Invasion Assay

Matrigel (Corning) was diluted to 300µg/ml in PBS and 100µl was added to an 8µm pore 24 well Boyden chamber insert (Corning). Plates were incubated at 37°C overnight to allow Matrigel to set. 25,000 cells per well were added in 500µl starvation media to the top of the chamber 750µl of complete media was added below the chamber. Cells were incubated for 24 hours at 37°C. Matrigel was poured from the chamber. Cells were fixed with 400µl of 4% PFA for 20 minutes, washed with PBS, permeabilised with 0.1% Triton x-100 in PBS for 15 minutes, washed with PBS and stained with 0.1% crystal violet (Sigma-Aldrich) in 4% PFA in PBS. Wells were washed with PBS and water and air dried. Absorbance at 590nm was used to assess crystal violet staining per well which shows the number of cells that migrated through the Matrigel plug and the insert membrane.

### Scratch Wound Cell Migration Assay

40,000 MDA-MB-231 cells and 70,000 MCF-7 cells per well were plated in a flat-bottomed 96 well plate (Corning) and incubated at 37°C overnight to reach 100% confluence. Cells were treated for 2 hours at 37°C with 10µg/ml mitomycin (Fisher Scientific) to prevent cell proliferation. An Essen bioscience woundmaker was used to create an even scratch in each well. Wells were washed with PBS and starvation media was added to each well. Recombinant OMD or CDK1 or CDK4,6 inhibitors were added to starvation media. Conditioned media was added in place of starvation media for conditioned media experiments. Images of each well were taken every hour using an Incucyte S3 system with 4x objective. Wound density at every hour over 48 hours was measured using ImageJ. The final wound density at 48 hours post-scratch was used as a measure of the degree of migration of cells.

### Mammosphere Assays

Flat-bottomed 6 well plates were coated with polyHEMA for 48 hours at 40°C until dry. Trypsinised cells were disaggregated to a single cell suspension by passing cells three times through a 25-gauge needle. 3000 disaggregated cells per well were plated into 2ml mammosphere media (DMEM/F12 with B27 supplement and 20ng/ml EGF (Sigma-Aldrich)) and cultured for 5 days in suspension before counting mammosphere greater than 50µm in size (width and height). Mammosphere forming efficiency as counted by dividing the number of mammosphere formed by the number of cells plated initially. Primary mammosphere were trypsin-digested and needle disaggregated as above and were plated at a density of 3000 cells per well into new polyHEMA-coated plates in 2ml of mammosphere media and incubated for 5 days in suspension. Mammosphere self-renewal percentage was calculated by dividing the number of secondary mammosphere that formed over the number of primary mammosphere and multiplying by 100.

### Alkaline Phosphatase Activity Assay

A p-Nitrophenyl Phosphate, Disodium Salt (PnPP) standard curve of 33µM, 67µM, 100µM, 133µM and 167µM in assay buffer was produced as a substrate curve for alkaline phosphatase modification. HFOB 1.19 cells were trypsinised, pelleted, resuspended in PBS and 50,000 cells were pelleted and 50µl of assay buffer and 50µl of lysis buffer (20mM Tris-HCl pH 8.0, 160mM NaCl, 1mM CaCl_2_, 1% Triton-X-100) and pipetted up and down 20 times to lyse cells. Samples were centrifuged at 18000xg for 15 minutes at 4°C to pellet insoluble material. 20µl of each supernatant was added to 60µl of assay buffer and 50µl 5mM PnPP in a 96 well plate. 10µl of alkaline phosphatase enzyme was added to each standard well and mixed by pipetting up and down. Samples were incubated at room temperature in the dark for 1 hour. 20µl of stop solution was added to each sample and optical density at 405nm was measured using a plate reader (Agilent/Biotek). Alkaline phosphatase activity was measured by comparing OD at 405nm to the standard curve of known alkaline phosphatase activity. Data were presented as µmol/min/ml of substrate modified by alkaline phosphatase per minute per ml of reaction mixture.

### Alizarin Red Staining

HFOB 1.19 cells were cultured at 34°C for undifferentiated cells or 39°C for differentiation. Media was changed every 2 days, including treatments such as recombinant OMD. Conditioned media was added 1:1 with HFOB 1.19 media. When cells reached 80% (for 34°C condition) or after 7 days (for 39°C condition) they were fixed with ethanol, washed with PBS and stained with 40mM Alizarin red pH 4.2 in water for 1 hour at room temperature on a rocker. Plates were washed with 95% ethanol until all unbound stain was removed. Plates were scanned on a GelCount plate imager (Oxford Optronix) and the percentage area stained with calcium per well was calculated using ImageJ.

### Bone Resorption Assay

Bovine cortical bone slices were UV-sterilised and added to a flat-bottomed 96 well plate. 50,000 RAW 264.7 cells (undifferentiated or RANKL-differentiated) were added to each well in 100µl media (complete media or conditioned media) including any DMSO or recombinant OMD treatment required. Cells were cultured for 7 days and media was changed every 48-72 hours. Bone discs were fixed in 4% PFA in PBS for 10 minutes at room temperature. Samples were stained with 1% toluidine blue in water for 4 minutes. Samples were sonicated at 25% amplitude (50-60Hz) for 30 minutes in deionised water to remove bound cells which may be blocking pits from being stained. Samples were stained again with 1% toludine blue in water for 4 minutes. Samples were washed with deionised water, air dried and imaged using a Leica M205 Stereo Fluorescence System at 0.912X magnification. Resorption pits were quantified and measured using ImageJ where pits of a minimum of 0.01mm in width and length were counted.

### Alamar Blue Cell Viability Assay

U-bottomed 96 well plates were coated with 100µl polyHEMA for 48 hours at 40°C until dry. 10,000 cells were plated in DMEM (for MDA-MB-231 cells) with 10% FCS, 1% Pen-Strep, 1mM Sodium Pyruvate or Advanced DMEM/F12 with FCS, 1ug/ml hydrocortisone and 20ng/ml EGF (for cells from metastatic effusion samples). 10µl alamar blue reagent was added to each well and incubated for 4 hours at 37°C. Fluorescence at 590nm was measured using a VarioSkan Lux with SkanIT 6.1 software. At least 3 biological replicates were performed for each condition for MDA-MB-231 cells and 3 technical replicates were performed for effusion samples from one effusion drain per patient.

### Quantification of OMD in conditioned medium and metastatic fluids

Conditioned medium (CM) was derived using serum free medium. 231-OMD, HMFU19 and 544R were cultured at 37°C and 5% CO_2_ for 72hrs. At 24-hour intervals, the medium was collected, pooled, and replaced with fresh serum free medium. The CM was then centrifuged at 500g for 20 minutes and then concentrated using Vivaspin Turbo 15 Centrifugal Concentrator 10 kDa PES tubes. The resulting samples and the metastatic fluids derived from ascitic drains and pleural effusions were then analysed using a Human Osteoadherin/Osteomodulin ELISA Kit following manufacturer’s instructions Briefly, 100 µL of concentrated medium from each condition was added and incubated for 2.5hrs. Following washing, 100 µL of biotinylated antibody was incubated for 1hr. The wells were washed again and 100 µL of Streptavidin was added and incubated for 45mins. Following washing, 100 µL of TMB One-Step Substrate Reagent was added and incubated in the dark for 30mins. Finally, 50 µL of stop solution was added and read at 450 nm.

### Statistics

Experiments were performed in at least three independent biological triplicates. One-way ANOVAs with Dunnett’s multiple comparisons test were used to determine significance between 3 or more experimental conditions. In experiments where the data was not normal, Kruskal-Wallis with Dunn’s multiple comparisons test was used to determine significance between 3 or more experimental conditions. Student’s t tests were used to determine significance between two experimental conditions.

## References

1. Nolan, E., Lindeman, G.J., and Visvader, J.E. (2023). Deciphering breast cancer: from biology to the clinic. Cell 186, 1708–1728. 10.1016/j.cell.2023.01.040.

2. Riggio, A.I., Varley, K.E., and Welm, A.L. (2021). The lingering mysteries of metastatic recurrence in breast cancer. British Journal of Cancer 124, 13–26. 10.1038/s41416-020-01161-4.

3. Fornetti, J., Welm, A.L., and Stewart, S.A. (2018). Understanding the Bone in Cancer Metastasis. J Bone Miner Res 33, 2099–2113. 10.1002/jbmr.3618.

4. Mundy, G.R. (2002). Metastasis to bone: causes, consequences and therapeutic opportunities. Nat Rev Cancer 2, 584–593. 10.1038/nrc867.

5. Satcher, R.L., and Zhang, X.H. (2022). Evolving cancer-niche interactions and therapeutic targets during bone metastasis. Nat Rev Cancer 22, 85–101. 10.1038/s41568-021-00406-5.

6. Zhang, W., Bado, I.L., Hu, J., Wan, Y.W., Wu, L., Wang, H., Gao, Y., Jeong, H.H., Xu, Z., Hao, X., et al. (2021). The bone microenvironment invigorates metastatic seeds for further dissemination. Cell 184, 2471–2486 e2420. 10.1016/j.cell.2021.03.011.

7. Coleman, R.E., and Rubens, R.D. (1987). The clinical course of bone metastases from breast cancer. Br J Cancer 55, 61–66. 10.1038/bjc.1987.13.

8. Parsons, J., and Francavilla, C. (2019). ’Omics Approaches to Explore the Breast Cancer Landscape. Front Cell Dev Biol 7, 395. 10.3389/fcell.2019.00395.

9. Paul, M.R., Pan, T.C., Pant, D.K., Shih, N.N., Chen, Y., Harvey, K.L., Solomon, A., Lieberman, D., Morrissette, J.J., Soucier-Ernst, D., et al. (2020). Genomic landscape of metastatic breast cancer identifies preferentially dysregulated pathways and targets. J Clin Invest 130, 4252–4265. 10.1172/jci129941.

10. Zhang, Y., He, W., and Zhang, S. (2019). Seeking for Correlative Genes and Signaling Pathways With Bone Metastasis From Breast Cancer by Integrated Analysis. Front Oncol 9, 138. 10.3389/fonc.2019.00138.

11. Krug, K., Jaehnig, E.J., Satpathy, S., Blumenberg, L., Karpova, A., Anurag, M., Miles, G., Mertins, P., Geffen, Y., Tang, L.C., et al. (2020). Proteogenomic Landscape of Breast Cancer Tumorigenesis and Targeted Therapy. Cell 183, 1436–1456.e1431. 10.1016/j.cell.2020.10.036.

12. Anurag, M., Jaehnig, E.J., Krug, K., Lei, J.T., Bergstrom, E.J., Kim, B.-J., Vashist, T.D., Tran Huynh, A.M., Dou, Y., Gou, X., et al. (2022). Proteogenomic markers of chemotherapy resistance and response in triple negative breast cancer. Cancer Discovery, CD-22-0200. 10.1158/2159-8290.CD-22-0200.

13. Zagorac, I., Fernandez-Gaitero, S., Penning, R., Post, H., Bueno, M.J., Mouron, S., Manso, L., Morente, M.M., Alonso, S., Serra, V., et al. (2018). In vivo phosphoproteomics reveals kinase activity profiles that predict treatment outcome in triple-negative breast cancer. Nature Communications 9, 3501. 10.1038/s41467-018-05742-z.

14. Satpathy, S., Jaehnig, E.J., Krug, K., Kim, B.-J., Saltzman, A.B., Chan, D.W., Holloway, K.R., Anurag, M., Huang, C., Singh, P., et al. (2020). Microscaled proteogenomic methods for precision oncology. Nature Communications 11, 532. 10.1038/s41467-020-14381-2.

15. Ramstad, V.E., Franzen, A., Heinegard, D., Wendel, M., and Reinholt, F.P. (2003). Ultrastructural distribution of osteoadherin in rat bone shows a pattern similar to that of bone sialoprotein. Calcif Tissue Int 72, 57–64. 10.1007/s00223-002-2047-9.

16. Sommarin, Y., Wendel, M., Shen, Z., Hellman, U., and Heinegard, D. (1998). Osteoadherin, a cell-binding keratan sulfate proteoglycan in bone, belongs to the family of leucine-rich repeat proteins of the extracellular matrix. J Biol Chem 273, 16723–16729. 10.1074/jbc.273.27.16723.

17. Watson, J., Ferguson, H.R., Brady, R.M., Ferguson, J., Fullwood, P., Mo, H., Bexley, K.H., Knight, D., Howell, G., Schwartz, J.M., et al. (2022). Spatially resolved phosphoproteomics reveals fibroblast growth factor receptor recycling-driven regulation of autophagy and survival. Nat Commun 13, 6589. 10.1038/s41467-022-34298-2.

18. Song, K., Yu, Z., Zu, X., Li, G., Hu, Z., and Xue, Y. (2022). Collagen Remodeling along Cancer Progression Providing a Novel Opportunity for Cancer Diagnosis and Treatment. Int J Mol Sci 23. 10.3390/ijms231810509.

19. Roudier, M.P., Winters, B.R., Coleman, I., Lam, H.M., Zhang, X., Coleman, R., Chéry, L., True, L.D., Higano, C.S., Montgomery, B., et al. (2016). Characterizing the molecular features of ERG-positive tumors in primary and castration resistant prostate cancer. Prostate 76, 810–822. 10.1002/pros.23171.

20. Kojima, K., Shimanuki, M., Shikami, M., Andreeff, M., and Nakakuma, H. (2009). Cyclin-dependent kinase 1 inhibitor RO-3306 enhances p53-mediated Bax activation and mitochondrial apoptosis in AML. Cancer Sci 100, 1128–1136. 10.1111/j.1349-7006.2009.01150.x.

21. O’Hare, M.J., Bond, J., Clarke, C., Takeuchi, Y., Atherton, A.J., Berry, C., Moody, J., Silver, A.R., Davies, D.C., Alsop, A.E., et al. (2001). Conditional immortalization of freshly isolated human mammary fibroblasts and endothelial cells. Proc Natl Acad Sci U S A 98, 646–651. 10.1073/pnas.98.2.646.

22. Harris, S.A., Enger, R.J., Riggs, L.B., and Spelsberg, T.C. (1995). Development and characterization of a conditionally immortalized human fetal osteoblastic cell line. Journal of Bone and Mineral Research 10, 178–186. 10.1002/jbmr.5650100203.

23. Lampiasi, N., Russo, R., Kireev, I., Strelkova, O., Zhironkina, O., and Zito, F. (2021). Osteoclasts Differentiation from Murine RAW 264.7 Cells Stimulated by RANKL: Timing and Behavior. Biology (Basel) 10. 10.3390/biology10020117.

24. Papadaki, V., Asada, K., Watson, J.K., Tamura, T., Leung, A., Hopkins, J., Dellett, M., Sasai, N., Davaapil, H., Nik-Zainal, S., et al. (2020). Two Secreted Proteoglycans, Activators of Urothelial Cell-Cell Adhesion, Negatively Contribute to Bladder Cancer Initiation and Progression. Cancers (Basel) 12. 10.3390/cancers12113362.

25. Macedo, F., Ladeira, K., Pinho, F., Saraiva, N., Bonito, N., Pinto, L., and Goncalves, F. (2017). Bone Metastases: An Overview. Oncol Rev 11, 321. 10.4081/oncol.2017.321.

26. Tashima, T., Nagatoishi, S., Sagara, H., Ohnuma, S., and Tsumoto, K. (2015). Osteomodulin regulates diameter and alters shape of collagen fibrils. Biochem Biophys Res Commun 463, 292–296. 10.1016/j.bbrc.2015.05.053.

27. Bos, P.D., Zhang, X.H., Nadal, C., Shu, W., Gomis, R.R., Nguyen, D.X., Minn, A.J., van de Vijver, M.J., Gerald, W.L., Foekens, J.A., and Massagué, J. (2009). Genes that mediate breast cancer metastasis to the brain. Nature 459, 1005–1009. 10.1038/nature08021.

28. Program, C.Z.I.C.S., Abdulla, S., Aevermann, B., Assis, P., Badajoz, S., Bell, S.M., Bezzi, E., Cakir, B., Chaffer, J., Chambers, S., et al. (2025). CZ CELLxGENE Discover: a single-cell data platform for scalable exploration, analysis and modeling of aggregated data. Nucleic Acids Res 53, D886–D900. 10.1093/nar/gkae1142.

29. Coleman, R., Cameron, D., Dodwell, D., Bell, R., Wilson, C., Rathbone, E., Keane, M., Gil, M., Burkinshaw, R., Grieve, R., et al. (2014). Adjuvant zoledronic acid in patients with early breast cancer: final efficacy analysis of the AZURE (BIG 01/04) randomised open-label phase 3 trial. Lancet Oncol 15, 997–1006. 10.1016/S1470-2045(14)70302-X.

30. Hornbeck, P.V., Kornhauser, J.M., Tkachev, S., Zhang, B., Skrzypek, E., Murray, B., Latham, V., and Sullivan, M. (2012). PhosphoSitePlus: a comprehensive resource for investigating the structure and function of experimentally determined post-translational modifications in man and mouse. Nucleic Acids Res 40, D261–270. 10.1093/nar/gkr1122.

31. Belguise, K., Milord, S., Galtier, F., Moquet-Torcy, G., Piechaczyk, M., and Chalbos, D. (2012). The PKCθ pathway participates in the aberrant accumulation of Fra-1 protein in invasive ER-negative breast cancer cells. Oncogene 31, 4889–4897. 10.1038/onc.2011.659.

32. Potapova, T.A., Daum, J.R., Byrd, K.S., and Gorbsky, G.J. (2009). Fine tuning the cell cycle: activation of the Cdk1 inhibitory phosphorylation pathway during mitotic exit. Mol Biol Cell 20, 1737–1748. 10.1091/mbc.e08-07-0771.

33. Mertins, P., Mani, D.R., Ruggles, K.V., Gillette, M.A., Clauser, K.R., Wang, P., Wang, X., Qiao, J.W., Cao, S., Petralia, F., et al. (2016). Proteogenomics connects somatic mutations to signalling in breast cancer. Nature 534, 55–62. 10.1038/nature18003.

34. Gyorffy, B. (2021). Survival analysis across the entire transcriptome identifies biomarkers with the highest prognostic power in breast cancer. Comput Struct Biotechnol J 19, 4101–4109. 10.1016/j.csbj.2021.07.014.

35. Vassilev, L.T., Tovar, C., Chen, S., Knezevic, D., Zhao, X., Sun, H., Heimbrook, D.C., and Chen, L. (2006). Selective small-molecule inhibitor reveals critical mitotic functions of human CDK1. Proc Natl Acad Sci U S A 103, 10660–10665. 10.1073/pnas.0600447103.

36. Rampersad, S.N. (2012). Multiple applications of Alamar Blue as an indicator of metabolic function and cellular health in cell viability bioassays. Sensors (Basel) 12, 12347–12360. 10.3390/s120912347.

37. Krek, W., and Nigg, E.A. (1991). Differential phosphorylation of vertebrate p34cdc2 kinase at the G1/S and G2/M transitions of the cell cycle: identification of major phosphorylation sites. EMBO J 10, 305–316. 10.1002/j.1460-2075.1991.tb07951.x.

38. Watase, C., Shiino, S., Shimoi, T., Noguchi, E., Kaneda, T., Yamamoto, Y., Yonemori, K., Takayama, S., and Suto, A. (2021). Breast Cancer Brain Metastasis-Overview of Disease State, Treatment Options and Future Perspectives. Cancers (Basel) 13. 10.3390/cancers13051078.

39. Kycler, W.L. P. (2011). Surgical approach to pulmonary metastases from breast cancer. The Breast Journal 18, 52–57.

40. Makhmut, A., Qin, D., Hartlmayr, D., Seth, A., and Coscia, F. (2024). An Automated and Fast Sample Preparation Workflow for Laser Microdissection Guided Ultrasensitive Proteomics. Mol Cell Proteomics 23, 100750. 10.1016/j.mcpro.2024.100750.

41. Petrosius, V., Aragon-Fernandez, P., Arrey, T.N., Woessman, J., Uresin, N., de Boer, B., Su, J., Furtwangler, B., Stewart, H., Denisov, E., et al. (2025). Quantitative Label-Free Single-Cell Proteomics on the Orbitrap Astral MS. Mol Cell Proteomics, 100982. 10.1016/j.mcpro.2025.100982.

42. Tashima, T., Nagatoishi, S., Caaveiro, J.M.M., Nakakido, M., Sagara, H., Kusano-Arai, O., Iwanari, H., Mimuro, H., Hamakubo, T., Ohnuma, S.I., and Tsumoto, K. (2018). Molecular basis for governing the morphology of type-I collagen fibrils by Osteomodulin. Commun Biol 1, 33 10.1038/s42003-018-0038-2.

43. Hamaya, E., Fujisawa, T., and Tamura, M. (2019). Osteoadherin serves roles in the regulation of apoptosis and growth in MC3T3-E1 osteoblast cells. Int J Mol Med 44, 2336–2344. 10.3892/ijmm.2019.4376.

44. Tan, Y., Demeter, M.R., Ruan, H., and Comb, M.J. (2000). BAD Ser-155 Phosphorylation Regulates BAD/Bcl-XL Interaction and Cell Survival *. Journal of Biological Chemistry 275, 25865–25869. 10.1074/jbc.M004199200.

45. Lin, W., Zhu, X., Gao, L., Mao, M., Gao, D., and Huang, Z. (2021). Osteomodulin positively regulates osteogenesis through interaction with BMP2. Cell Death Dis 12, 147. 10.1038/s41419-021-03404-5.

46. Giorello, M.B., Martinez, L.M., Borzone, F.R., Padin, M.D.R., Mora, M.F., Sevic, I., Alaniz, L., Calcagno, M.L., Garcia-Rivello, H., Wernicke, A., et al. (2023). CD105 expression in cancer-associated fibroblasts: a biomarker for bone metastasis in early invasive ductal breast cancer patients. Front Cell Dev Biol 11, 1250869. 10.3389/fcell.2023.1250869.

47. Katsuno, Y., Hanyu, A., Kanda, H., Ishikawa, Y., Akiyama, F., Iwase, T., Ogata, E., Ehata, S., Miyazono, K., and Imamura, T. (2008). Bone morphogenetic protein signaling enhances invasion and bone metastasis of breast cancer cells through Smad pathway. Oncogene 27, 6322–6333. 10.1038/onc.2008.232.

48. Sloan, E.K., Pouliot, N., Stanley, K.L., Chia, J., Moseley, J.M., Hards, D.K., and Anderson, R.L. (2006). Tumor-specific expression of alphavbeta3 integrin promotes spontaneous metastasis of breast cancer to bone. Breast Cancer Res 8, R20. 10.1186/bcr1398.

49. Mirzakhanyan, Y., Jankevics, A., Scheltema, R.A., and Gershon, P.D. (2023). Combination of deep XLMS with deep learning reveals an ordered rearrangement and assembly of a major protein component of the vaccinia virion. mBio 14, e0113523. 10.1128/mbio.01135-23.

50. Muller, M., Grabnitz, F., Barandun, N., Shen, Y., Wendt, F., Steiner, S.N., Severin, Y., Vetterli, S.U., Mondal, M., Prudent, J.R., et al. (2021). Light-mediated discovery of surfaceome nanoscale organization and intercellular receptor interaction networks. Nat Commun 12, 7036. 10.1038/s41467-021-27280-x.

51. Ninomiya, K., Miyamoto, T., Imai, J., Fujita, N., Suzuki, T., Iwasaki, R., Yagi, M., Watanabe, S., Toyama, Y., and Suda, T. (2007). Osteoclastic activity induces osteomodulin expression in osteoblasts. Biochem Biophys Res Commun 362, 460–466. 10.1016/j.bbrc.2007.07.193.

52. Rehn, A.P., Cerny, R., Sugars, R.V., Kaukua, N., and Wendel, M. (2008). Osteoadherin is upregulated by mature osteoblasts and enhances their in vitro differentiation and mineralization. Calcif Tissue Int 82, 454–464. 10.1007/s00223-008-9138-1.

53. Qiao, M., Shapiro, P., Fosbrink, M., Rus, H., Kumar, R., and Passaniti, A. (2006). Cell cycle-dependent phosphorylation of the RUNX2 transcription factor by cdc2 regulates endothelial cell proliferation. J Biol Chem 281, 7118–7128. 10.1074/jbc.M508162200.

54. Rajgopal, A., Young, D.W., Mujeeb, K.A., Stein, J.L., Lian, J.B., van Wijnen, A.J., and Stein, G.S. (2007). Mitotic control of RUNX2 phosphorylation by both CDK1/cyclin B kinase and PP1/PP2A phosphatase in osteoblastic cells. J Cell Biochem 100, 1509–1517. 10.1002/jcb.21137.

55. Li, X.Q., Lu, J.T., Tan, C.C., Wang, Q.S., and Feng, Y.M. (2016). RUNX2 promotes breast cancer bone metastasis by increasing integrin alpha5-mediated colonization. Cancer Lett 380, 78–86. 10.1016/j.canlet.2016.06.007.

56. Ran, R., Harrison, H., Syamimi Ariffin, N., Ayub, R., Pegg, H.J., Deng, W., Mastro, A., Ottewell, P.D., Mason, S.M., Blyth, K., et al. (2020). A role for CBFbeta in maintaining the metastatic phenotype of breast cancer cells. Oncogene 39, 2624–2637. 10.1038/s41388-020-1170-2.

57. Jones, M.C., Askari, J.A., Humphries, J.D., and Humphries, M.J. (2018). Cell adhesion is regulated by CDK1 during the cell cycle. Journal of Cell Biology 217, 3203–3218. 10.1083/jcb.201802088.

58. Ratnayake, W.S., Apostolatos, C.A., Breedy, S., Dennison, C.L., Hill, R., and Acevedo-Duncan, M. (2021). Atypical PKCs activate Vimentin to facilitate prostate cancer cell motility and invasion. Cell Adh Migr 15, 37–57. 10.1080/19336918.2021.1882782.

59. Li, M., He, F., Zhang, Z., Xiang, Z., and Hu, D. (2020). CDK1 serves as a potential prognostic biomarker and target for lung cancer. J Int Med Res 48, 300060519897508. 10.1177/0300060519897508.

60. Ravindran Menon, D., Luo, Y., Arcaroli, J.J., Liu, S., KrishnanKutty, L.N., Osborne, D.G., Li, Y., Samson, J.M., Bagby, S., Tan, A.C., et al. (2018). CDK1 Interacts with Sox2 and Promotes Tumor Initiation in Human Melanoma. Cancer Res 78, 6561–6574. 10.1158/0008-5472.Can-18-0330.

61. Zheng, H.P., Huang, Z.G., He, R.Q., Lu, H.P., Dang, Y.W., Lin, P., Wen, D.Y., Qin, Y.Y., Luo, B., Li, X.J., et al. (2019). Integrated assessment of CDK1 upregulation in thyroid cancer. Am J Transl Res 11, 7233–7254.

62. Wu, C.X., Wang, X.Q., Chok, S.H., Man, K., Tsang, S.H.Y., Chan, A.C.Y., Ma, K.W., Xia, W., and Cheung, T.T. (2018). Blocking CDK1/PDK1/β-Catenin signaling by CDK1 inhibitor RO3306 increased the efficacy of sorafenib treatment by targeting cancer stem cells in a preclinical model of hepatocellular carcinoma. Theranostics 8, 3737–3750. 10.7150/thno.25487.

63. Paraghamian, S.E., Huang, Y., Hawkins, G.M., Fan, Y., Yin, Y., Zhang, X., Suo, H., Zhou, C., and Bae-Jump, V.L. (2020). The Cdk1 inhibitor RO3306 has anti-tumorigenic effects in high-grade serous ovarian cancer. Gynecologic Oncology 159, 88–89. 10.1016/j.ygyno.2020.05.068.

64. Wang, Q., Bode, A.M., and Zhang, T. (2023). Targeting CDK1 in cancer: mechanisms and implications. NPJ Precis Oncol 7, 58. 10.1038/s41698-023-00407-7.

65. Ji, H., Wang, J., Nika, H., Hawke, D., Keezer, S., Ge, Q., Fang, B., Fang, X., Fang, D., Litchfield, D.W., et al. (2009). EGF-induced ERK activation promotes CK2-mediated disassociation of alpha-Catenin from beta-Catenin and transactivation of beta-Catenin. Mol Cell 36, 547–559. 10.1016/j.molcel.2009.09.034.

66. Kang, Y., Siegel, P.M., Shu, W., Drobnjak, M., Kakonen, S.M., Cordón-Cardo, C., Guise, T.A., and Massagué, J. (2003). A multigenic program mediating breast cancer metastasis to bone. Cancer Cell 3, 537–549. 10.1016/S1535-6108(03)00132-6.

67. Holliday, D.L., Brouilette, K.T., Markert, A., Gordon, L.A., and Jones, J.L. (2009). Novel multicellular organotypic models of normal and malignant breast: tools for dissecting the role of the microenvironment in breast cancer progression. Breast Cancer Res 11, R3. 10.1186/bcr2218.

68. Kojima, Y., Acar, A., Eaton, E.N., Mellody, K.T., Scheel, C., Ben-Porath, I., Onder, T.T., Wang, Z.C., Richardson, A.L., Weinberg, R.A., and Orimo, A. (2010). Autocrine TGF-beta and stromal cell-derived factor-1 (SDF-1) signaling drives the evolution of tumor-promoting mammary stromal myofibroblasts. Proc Natl Acad Sci U S A 107, 20009–20014. 10.1073/pnas.1013805107.

69. Collin-Osdoby, P., and Osdoby, P. (2012). RANKL-Mediated Osteoclast Formation from Murine RAW 264.7 cells. In Bone Research Protocols, M.H. Helfrich, and S.H. Ralston, eds. (Humana Press), pp. 187–202. 10.1007/978-1-61779-415-5_13.

70. Wilcock, D.J., Badrock, A.P., Wong, C.W., Owen, R., Guerin, M., Southam, A.D., Johnston, H., Telfer, B.A., Fullwood, P., Watson, J., et al. (2022). Oxidative stress from DGAT1 oncoprotein inhibition in melanoma suppresses tumor growth when ROS defenses are also breached. Cell Rep 39, 110995. 10.1016/j.celrep.2022.110995.

71. Cox, J., and Mann, M. (2008). MaxQuant enables high peptide identification rates, individualized p.p.b.-range mass accuracies and proteome-wide protein quantification. Nat Biotechnol 26, 1367–1372. 10.1038/nbt.1511.

72. Tyanova, S., Temu, T., Sinitcyn, P., Carlson, A., Hein, M.Y., Geiger, T., Mann, M., and Cox, J. (2016). The Perseus computational platform for comprehensive analysis of (prote)omics data. Nat Methods 13, 731–740. 10.1038/nmeth.3901.

73. Szklarczyk, D., Gable, A.L., Lyon, D., Junge, A., Wyder, S., Huerta-Cepas, J., Simonovic, M., Doncheva, N.T., Morris, J.H., Bork, P., et al. (2019). STRING v11: protein-protein association networks with increased coverage, supporting functional discovery in genome-wide experimental datasets. Nucleic Acids Res 47, D607–D613. 10.1093/nar/gky1131.

74. Kohl, M., Wiese, S., and Warscheid, B. (2011). Cytoscape: software for visualization and analysis of biological networks. Methods Mol Biol 696, 291–303. 10.1007/978-1-60761-987-1_18.

75. Schneider, C.A., Rasband, W.S., and Eliceiri, K.W. (2012). NIH Image to ImageJ: 25 years of image analysis. Nat Methods 9, 671–675. 10.1038/nmeth.2089.

76. Bankhead, P., Loughrey, M.B., Fernandez, J.A., Dombrowski, Y., McArt, D.G., Dunne, P.D., McQuaid, S., Gray, R.T., Murray, L.J., Coleman, H.G., et al. (2017). QuPath: Open source software for digital pathology image analysis. Sci Rep 7, 16878. 10.1038/s41598-017-17204-5.

77. Ritchie, M.E., Phipson, B., Wu, D., Hu, Y., Law, C.W., Shi, W., and Smyth, G.K. (2015). limma powers differential expression analyses for RNA-sequencing and microarray studies. Nucleic Acids Res 43, e47. 10.1093/nar/gkv007.

78. Wu, S.Z., Al-Eryani, G., Roden, D.L., Junankar, S., Harvey, K., Andersson, A., Thennavan, A., Wang, C., Torpy, J.R., Bartonicek, N., et al. (2021). A single-cell and spatially resolved atlas of human breast cancers. Nat Genet 53, 1334–1347. 10.1038/s41588-021-00911-1.

