## Supplemental Information for "Proteomics Identifies Osteomodulin as a Promoter of Breast Cancer Bone Metastasis via CDK1 Activation"

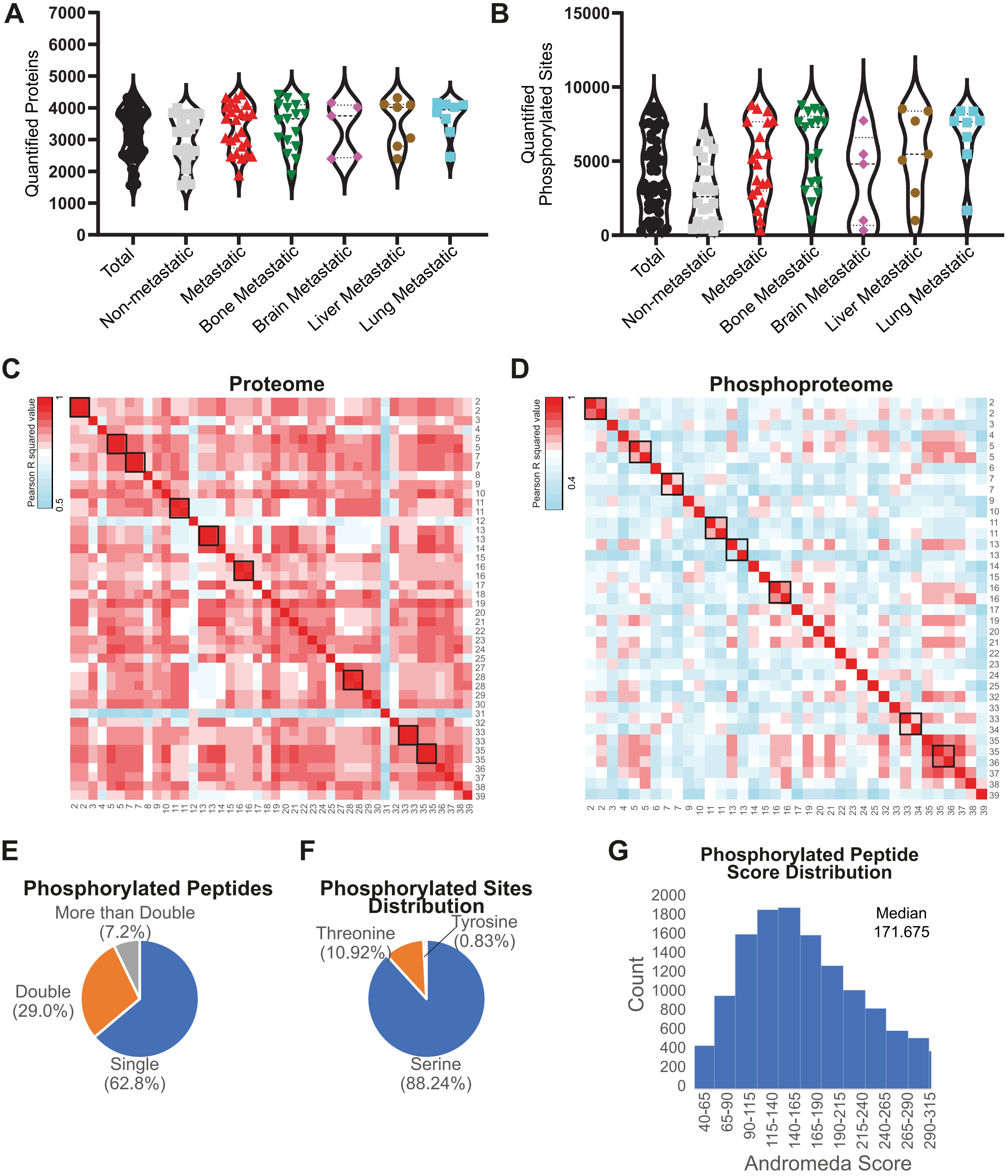
A
Quantified Proteins
7000
6000
5000
4000
3000
2000
1000
0
Total
Non-metastatic
Metastatic
Bone Metastatic
Brain Metastatic
Liver Metastatic
Lung Metastatic
B
Quantified Phosphorylated Sites
15000
10000
5000
0
Total
Non-metastatic
Metastatic
Bone Metastatic
Brain Metastatic
Liver Metastatic
Lung Metastatic
C
Proteome
Pearson R squared value
1
0.5
D
Phosphoproteome
Pearson R squared value
1
0.4
E
Phosphorylated Peptides
More than Double (7.2%)
Double (29.0%)
Single (62.8%)
F
Phosphorylated Sites Distribution
Threonine (10.92%)
Tyrosine (0.83%)
Serine (88.24%)
G
Phosphorylated Peptide Score Distribution
Count
2000
1800
1600
1400
1200
1000
800
600
400
200
0
Median
171.675
40-65
65-90
90-115
115-140
140-165
165-190
190-215
215-240
240-265
265-290
290-315
Andromeda Score

**Figure S1.** Proteomics and phosphoproteomics analysis of metastatic and non-metastatic samples shows good quality of the data. **A, B,** Distribution of the number of proteins (A) and phosphorylated sites (B) identified and quantified in at least one patient (total), one patient who did not develop metastases within 5 years of diagnosis (non-metastatic), one patient who did develop metastases within 5 years of diagnosis (metastatic), and in patients who developed bone, brain, liver or lung metastases within 5 years of diagnosis. **C, D,** Heatmap showing the Pearson correlation of patient samples in the proteome (A) and phosphoproteome (B). Black squares indicate technical replicates starting from materials from the same patient. **E,** Distribution of phosphorylated peptides with one, two or more phosphorylated sites. **F,** Distribution of identified serine, threonine, and tyrosine phosphorylated sites. **G,** Distribution of phosphorylated peptide Andromeda scores

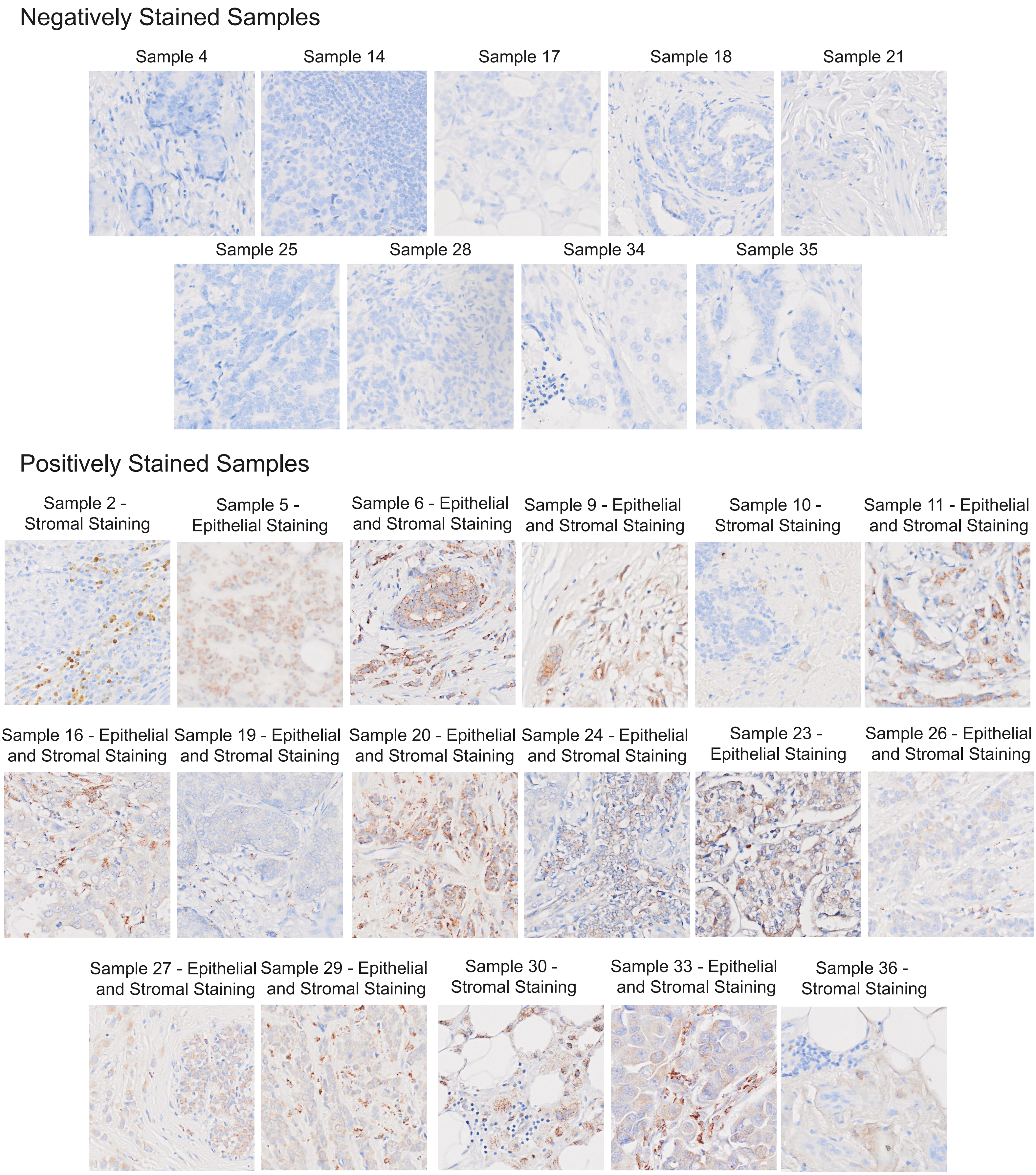
Negatively Stained Samples
Sample 4
Sample 14
Sample 17
Sample 18
Sample 21
Sample 25
Sample 28
Sample 34
Sample 35
Positively Stained Samples
Sample 2 - Stromal Staining
Sample 5 - Epithelial Staining
Sample 6 - Epithelial and Stromal Staining
Sample 9 - Epithelial and Stromal Staining
Sample 10 - Stromal Staining
Sample 11 - Epithelial and Stromal Staining
Sample 16 - Epithelial and Stromal Staining
Sample 19 - Epithelial and Stromal Staining
Sample 20 - Epithelial and Stromal Staining
Sample 24 - Epithelial and Stromal Staining
Sample 23 - Epithelial Staining
Sample 26 - Epithelial and Stromal Staining
Sample 27 - Epithelial and Stromal Staining
Sample 29 - Epithelial and Stromal Staining
Sample 30 - Stromal Staining
Sample 33 - Epithelial and Stromal Staining
Sample 36 - Stromal Staining

**Figure S2.** Immunohistochemical staining analysis shows OMD expression in epithelial and stromal compartments of patient-derived samples. Immunohistochemical staining of the 26 patient-derived samples shows no staining in 9 samples and the localization of OMD in epithelial and stromal compartments in 17 samples.

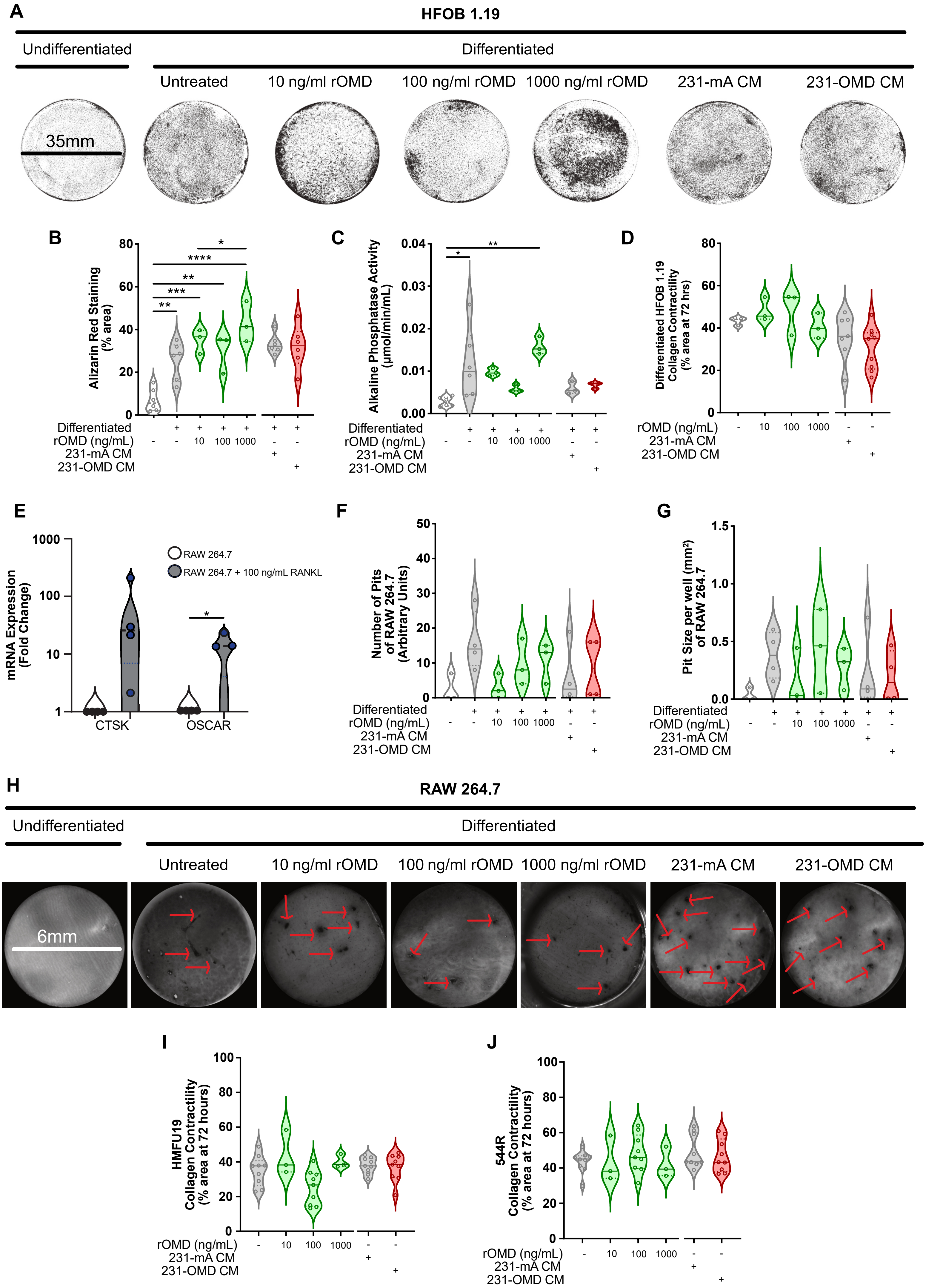
A
HFOB 1.19
Undifferentiated
Differentiated
Untreated
10 ng/ml rOMD
100 ng/ml rOMD
1000 ng/ml rOMD
231-mA CM
231-OMD CM
35mm
B
Alizarin Red Staining (% area)
Differentiated
rOMD (ng/mL)
231-mA CM
231-OMD CM
C
Alkaline Phosphatase Activity (µmol/min/mL)
D
Differentiated HFOB 1.19 Collagen Contractility (% area at 72 hrs)
E
mRNA Expression (Fold Change)
RAW 264.7
RAW 264.7 + 100 ng/mL RANKL
CTSK
OSCAR
F
Number of Pits of RAW 264.7 (Arbitrary Units)
G
Pit Size per well (mm²) of RAW 264.7
H
RAW 264.7
6mm
I
HMFU19 Collagen Contractility (% area at 72 hours)
J
544R Collagen Contractility (% area at 72 hours)

**Figure S3.** OMD does not affect the function of osteoclasts and fibroblasts and partially affects the function of osteoblasts present within the microenvironment. **A,** Representative images from N= 3 independent biological replicates of HFOB 1.19 osteoblast cells left undifferentiated or differentiated by growth at 39.5°C for 7 days, treated (or not) with the indicated concentrations of rOMD or with conditioned medium from either 231-mA or 231-OMD, and subjected to alizarin red staining of calcium deposition. Black areas indicate calcium staining. Scale bar, 35mm. **B, C,** Alizarin red staining (B) and alkaline phosphatase activity (C) of hFOB 1.19 osteoblasts grown at 34°C (undifferentiated) or 39.5°C (differentiated) followed by treatment (or not) with conditioned media or rOMD. **D,** Relative mRNA expression of Cathepsin K (CTSK) and OSCAR from RAW 264.7 cells differentiated with 100ng/ml RANKL for 8 days compared to untreated RAW 264.7 osteoclast cells and assessed by qPCR. Data is presented as the ratio between treated and control from N= 3 independent biological replicates. (mean 65.65 and 12.95 respectively) p = <0.05 *, <0.01 **, <0.001 ***, <0.0001 **** (Student's t test). **E, F,** Bone resorption assay assessing pit formation (F) and size of pits per well/bone disc (G) of RAW 264.7 monocyte cells differentiated or not to osteoclasts with RANKL and treated or not with conditioned media or rOMD. Data is presented as mean +/- SEM of N = 3 independent biological replicates. Experiments with CM were performed independently of rOMD experiments and were thus not directly compared. p = <0.05 *, <0.01 **, <0.001 ***, <0.0001 **** (One Way ANOVA with Dunnett's Multiple Comparison Test, compared to untreated control). **G,** Representative images from N= >3 independent biological replicates of bone resorption assay where bovine bone discs were cultured with RAW 264.7 either left undifferentiated or differentiated with RANKL as in B and treated or not with the indicated concentrations of rOMD and with OMD conditioned media from 231-mA and 231-OMD. Red arrows points to black dots indicating pores formed by osteoclast bone resorption. Scale bar, 6mm. **H, I, J,** Collagen contractility assay of differentiated HFOB 1.19 osteoblast cells, HMFU19 immortalised normal fibroblasts (I) and 544R immortalised cancer-associated fibroblasts (J) treated with conditioned medium (CM) from either 231-mA or 231-OMD and the indicated concentrations of rOMD where collagen plug shrinkage to a % area of its original size over 72 hours represents collagen remodelling compared to time.

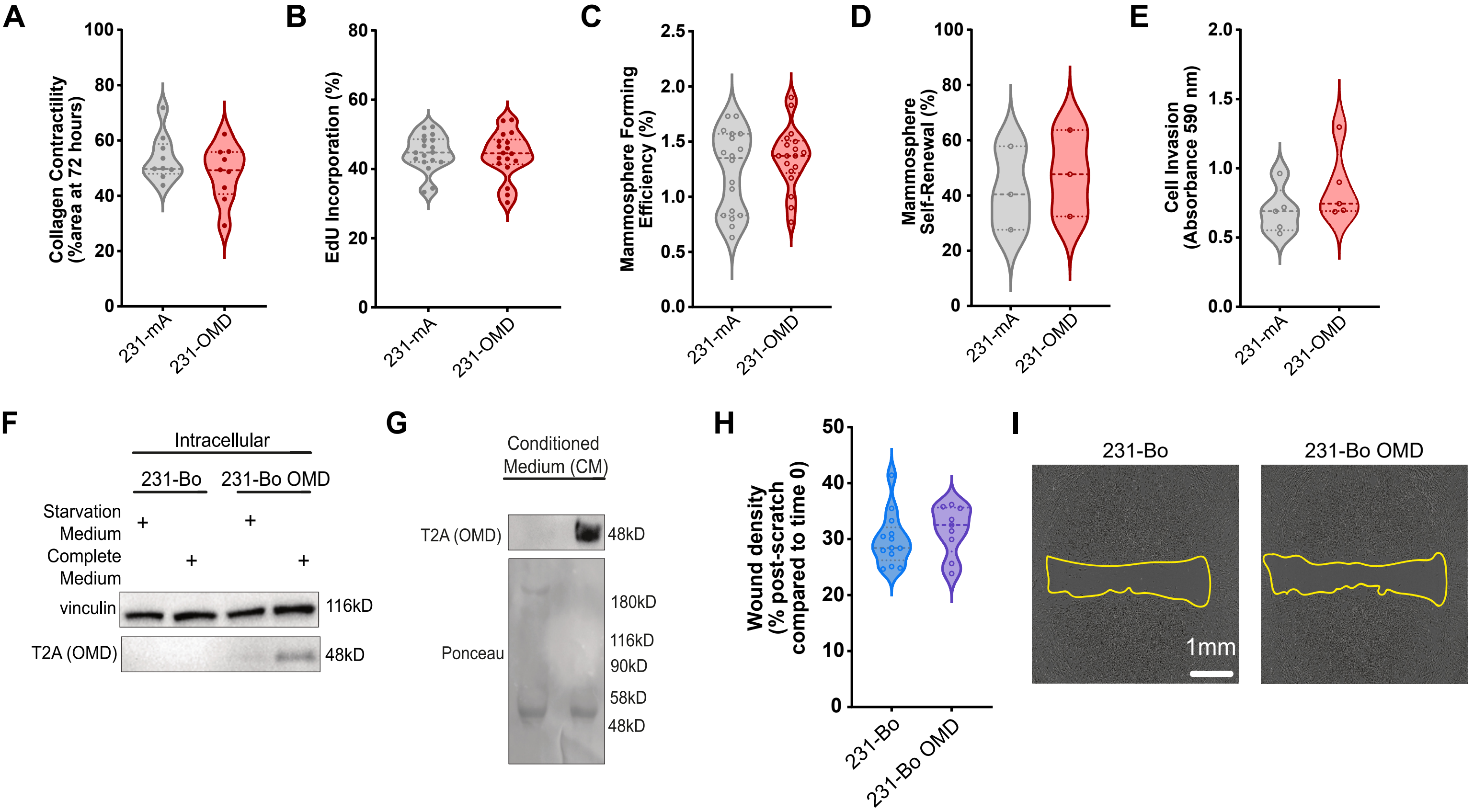
A
Collagen Contractility (%area at 72 hours)
231-mA
231-OMD
B
EdU Incorporation (%)
C
Mammosphere Forming Efficiency (%)
D
Mammosphere Self-Renewal (%)
E
Cell Invasion (Absorbance 590 nm)
F
Intracellular
231-Bo
231-Bo OMD
Starvation Medium
Complete Medium
vinculin
116kD
T2A (OMD)
48kD
G
Conditioned Medium (CM)
T2A (OMD)
48kD
Ponceau
180kD
116kD
90kD
58kD
48kD
H
Wound density (% post-scratch compared to time 0)
231-Bo
231-Bo OMD
I
231-Bo
231-Bo OMD
1mm

**Figure S4.** OMD does not affect other metastatic behaviours of breast cancer cells. **A,** Collagen contractility assay of 231-mA and 231-OMD cells where collagen plug shrinkage to a % area of its original size over 72 hours represents collagen remodelling compared to time zero. **B,** Percentage of EdU incorporation of 231-mA and 231-OMD cells at 4h compared to time zero. **C,** Mammosphere forming efficiency of 231-mA and 231-OMD cells after 5 days in non-adherent culture. **D,** Mammosphere self-renewal of disaggregated primary mammosphere grown in non-adherent culture for 5 days from C. **E,** Boyden chamber invasion assay through Matrigel measured by absorbance of crystal violet-stained cells at 590nm after 24h. **F, G,** Immunoblotting analysis with the indicated antibodies of total lysates from the MDA-MB-231 bone tropic variant cells (24) stably transfected with mApple (231-Bo) or an OMD-T2A-mApple construct (231-Bo OMD) and grown in complete or starved medium. and of conditioned medium from the same cells. N = 3 independent biological replicates. **H,** Cell migration of 231-Bo (blue) and 231-Bo OMD (purple) assessed by wound density (percentage of the wound filled with cells 48hrs post-scratch compared to time zero). Data represents the mean +/- SEM of N = 3 independent biological replicates, each including at least 3 technical triplicates. **I,** Representative images from G. Scale bar = 1mm. **A-E,** Data is presented as mean +/- SEM of N = at least 3 independent biological experiments. *P* =< 0.05*, < 0.01**, < 0.001***, < 0.0001**** Mean +/- SEM (Student's *t*-test).

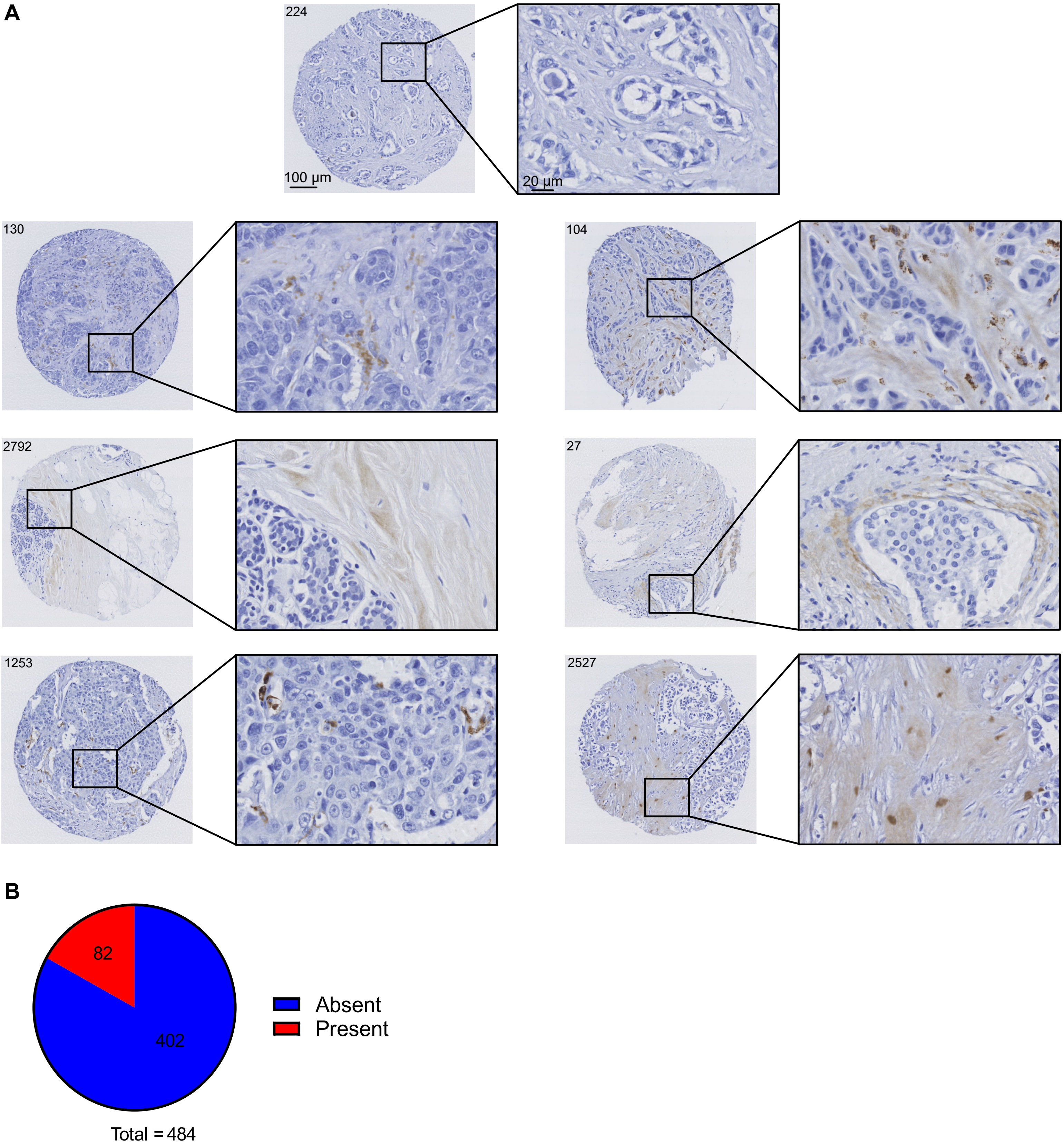
A
224
100 µm
20 µm
130
104
2792
27
1253
2527
B
82
402
Absent
Present
Total = 484

**Figure S5.** Immunohistochemical staining analysis shows OMD expression in stromal compartments of patient-derived samples from the AZURE trial. **A,** Expression of OMD in the stroma from the primary tumour of breast cancer patients. **B,** Proportion of patients that stained positive (red) with OMD and that stained negative (blue). (N = 484 patients)

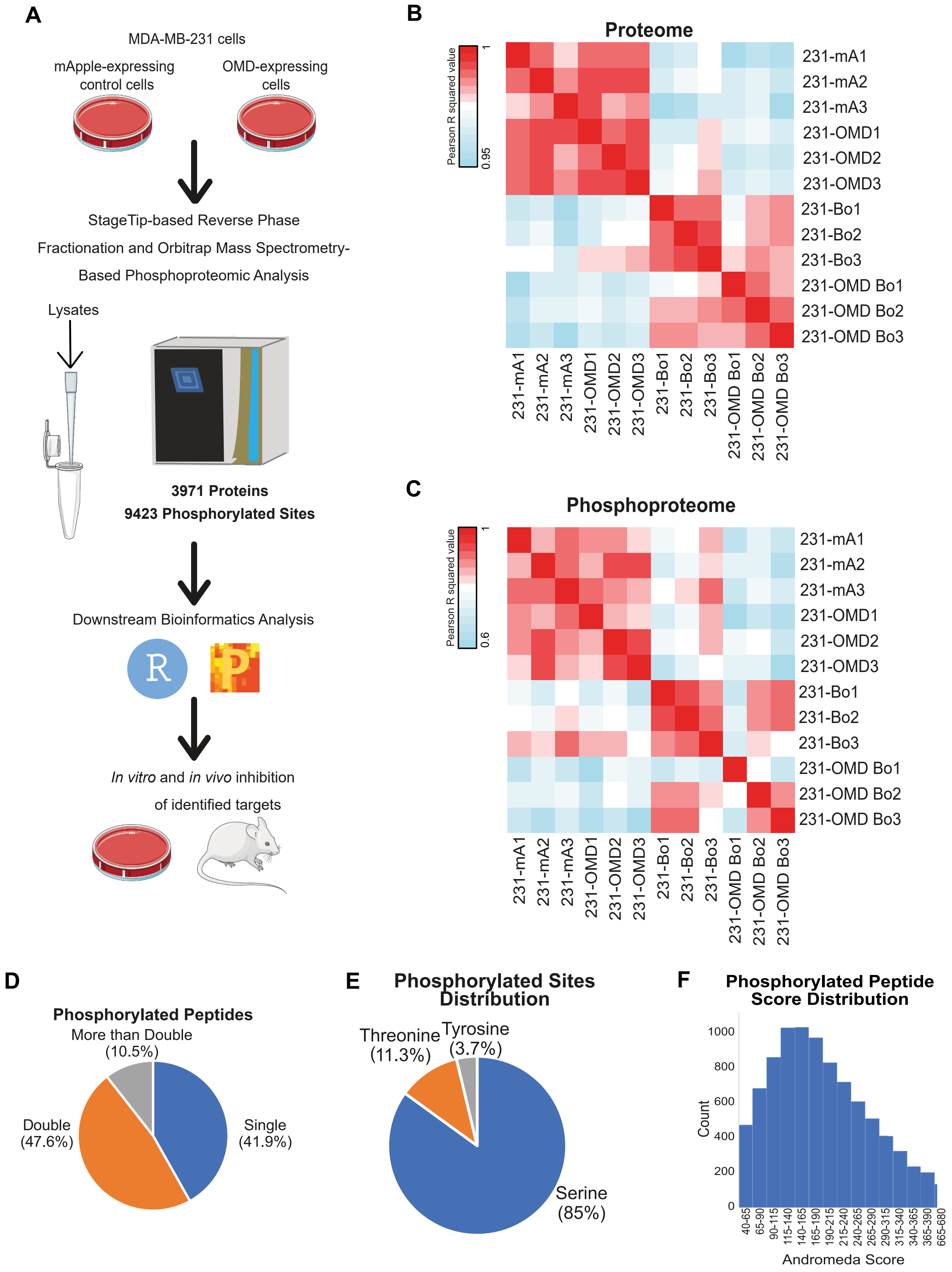

A
MDA-MB-231 cells
mApple-expressing control cells
OMD-expressing cells
StageTip-based Reverse Phase Fractionation and Orbitrap Mass Spectrometry-Based Phosphoproteomic Analysis
Lysates
3971 Proteins
9423 Phosphorylated Sites
Downstream Bioinformatics Analysis
R
*In vitro* and *in vivo* inhibition of identified targets
B
Proteome
Pearson R squared value
1
0.95
231-mA1
231-mA2
231-mA3
231-OMD1
231-OMD2
231-OMD3
231-Bo1
231-Bo2
231-Bo3
231-OMD Bo1
231-OMD Bo2
231-OMD Bo3
C
Phosphoproteome
Pearson R squared value
1
0.6
D
Phosphorylated Peptides
More than Double (10.5%)
Double (47.6%)
Single (41.9%)
E
Phosphorylated Sites Distribution
Threonine (11.3%)
Tyrosine (3.7%)
Serine (85%)
F
Phosphorylated Peptide Score Distribution
Count
1000
800
600
400
200
0
40-65
65-90
90-115
115-140
140-165
165-190
190-215
215-240
240-265
265-290
290-315
315-340
340-365
365-390
665-680
Andromeda Score

**Figure S6.** Proteomics and phosphoproteomics analysis of 231-mA, 231-OMD, 231-Bo, 231-Bo OMD cells and patient-derived metastatic effusion samples show good quality. **A,** Workflow of the proteomics and phosphoproteomics analysis of 231-mA, 231-OMD, 231-Bo, 231-Bo OMD cells. The numbers of proteins and phosphorylated sites identified are indicated. N = 3 Independent biological replicates. Tissue and mouse images were downloaded from SmartServier Medical Art. **B, C,** Heatmap showing the Pearson correlation of patient samples in the proteome (B) and phosphoproteome (C). **D,** Distribution of phosphorylated peptides with one, two or more phosphorylated sites. **E,** Distribution of identified serine, threonine, and tyrosine phosphorylated sites. **F,** Distribution of phosphorylated peptide Andromeda scores.

**A**

| Sample ID | Sample Type | Tumour pathology | | | Metastatic Sample Pathology | | | | | Sites of metastases at time of collection |
|---|---|---|---|---|---|---|---|---|---|---|
| | | ER | PR | HER2 | %ER | %PR | HER2 | Epithelial (% CK18) | Lymphocyte (% CD45) | |
| BB3RC29 | Ascitic Drain | + | + | - | 0 | 0 | 0 | 96 | 3 | Bone (Lytic and Sclerotic), Pleural, Retroperitoneal Lymph Nodes |
| BB3RC44 | Ascitic Drain | + | + | - | 24 | 0 | 0 | 90 | 4 | Peritoneum, Serosa |
| BB3RC45 | Ascitic Drain | + | + | - | 0 | 0 | 0 | 94 | 4 | No Information Available |
| BB3RC61 | Ascitic Drain | + | + | - | 0 | 0 | 0 | 87 | 12 | Peritoneum, Serosa |
| BB3RC66 | Ascitic Drain | + | + | - | 0 | 0 | 0 | 97 | 1 | Bone (Sclerotic), Peritoneum, Serosa |
| BB3RC71 | Ascitic Drain | + | + | + | 96 | 2 | 0 | 90 | 8 | Liver, Bone |
| BB3RC79 | Pleural Effusion | - | - | - | 0 | 0 | 0 | 58 | 25 | Lung |
| BB7042 | Pleural Effusion | + | + | + | 32 | 100 | 100 | 38 | 59 | Bone, Liver, Lung |

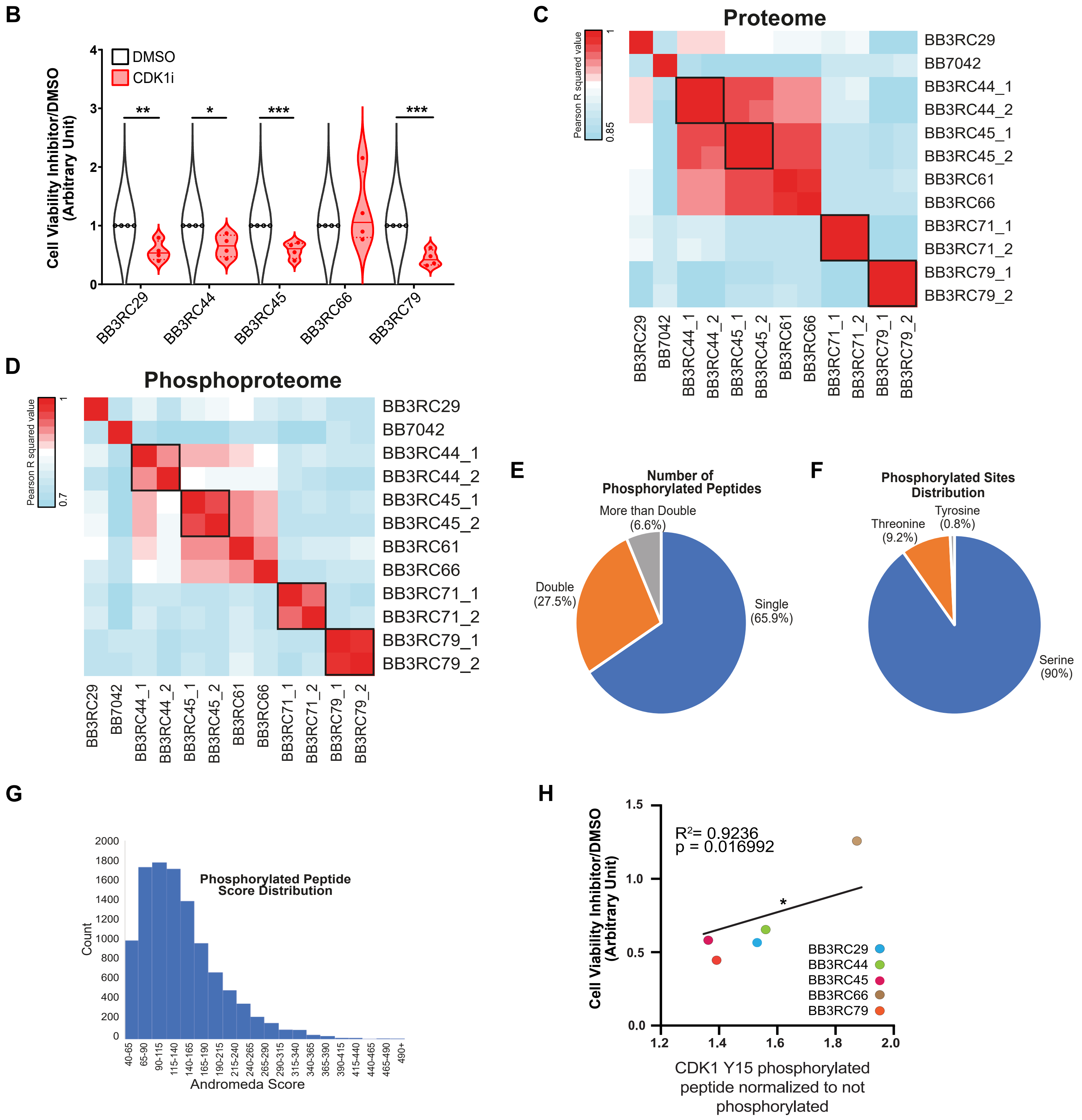

**Figure S7.** Proteomics and phosphoproteomics analysis of patient-derived metastatic effusion samples correlates CDK1 phosphorylation status with viability in response to CDK1 inhibition. **A,** Clinical and pathological characteristics of fresh frozen metastatic effusion samples. **B,** Relative viability of 5 metastatic pleural effusion measured by cell fluorescence at 590nm upon reduction of Alamar blue stain during incubation with non-adherent cells for 4 hours. Data is presented as mean +/- SEM of N >=3 independent biological replicates. P =< 0.05*, < 0.01**, < 0.001***, < 0.0001**** (Student's t-test). **C, D,** Heatmap showing the Pearson correlation of the proteome (C) and phosphoproteome of patient-derived metastatic effusion samples (D). Black squares indicate technical replicates starting from materials from the same patient based on Tables S13 and S14. **E,** Distribution of phosphorylated peptides with one, two or more phosphorylated sites. **F,** Distribution of identified serine, threonine, and tyrosine phosphorylated sites. **G,** Distribution of phosphorylated peptide Andromeda scores. **G, H,** Correlation between CDK1 Y15 normalised over CDK1 total protein normalised intensity and viability in response to CDK1 inhibition with 10μM RO-3306. Data is presented as mean response to viability, N=4 independent biological replicates. P =< 0.05*, < 0.01**, < 0.001***, < 0.0001**** (One-sided Linear Regression t test).

**Supplemental Information titles and legends**

Table S1. Demographic and clinical characteristics of fresh frozen primary breast cancer biopsy samples.

Table S2. Quality control-filtered proteome analysis of fresh frozen primary breast cancer biopsy samples.

Table S3. Quality control-filtered phosphoproteome analysis of fresh frozen primary breast cancer biopsy samples.

Table S4. Differential expression analysis of proteome data comparing patients who developed metastases within 5 years of diagnosis to those who did not develop metastases.

Table S5. Differential expression analysis of phosphoproteome data comparing patients who developed metastases within 5 years of diagnosis to those who did not develop metastases.

Table S6. A literature-curated database of genes, RNA transcripts, proteins and phosphorylated sites associated with the positive or negative regulation of breast cancer metastasis.

Table S7. KEGG and GO enrichment analysis of proteins and phosphorylated sites differentially regulated between patients who developed metastases within 5 years of diagnosis to those who did not develop metastases.

Table S8. Quality control-filtered proteome analysis of 231-mA, 231-OMD and 231-Bo OMD.

Table S9. Quality control-filtered phosphoproteome analysis of 231-mA, 231-OMD and 231-Bo OMD.

Table S10. Differential expression analysis of proteome data comparing 231-mA, 231-OMD and 231-Bo OMD.

Table S11. Differential expression analysis of phosphoproteome data comparing 231-mA and 231-OMD or 231-Bo OMD.

Table S12. Clinicopathological characteristics of patients used for assessment of independent prognostic value of OMD and CDK1.

Table S13. Quality control-filtered proteome analysis of cells from breast cancer metastatic effusions.

Table S14. Quality control-filtered phosphoproteome analysis of cells from breast cancer metastatic effusions.
